# The impact of placental structure on haemodynamics and fetal oxygen uptake

**DOI:** 10.64898/2026.02.02.703237

**Authors:** Z. Crowson, A. Blakey, D. Amanitis, R. Mcnair, L. Leach, C.A. Whitfield, I.L. Chernyavsky, O.E. Jensen, P. Houston, M.E. Hubbard, R.D. O’Dea

## Abstract

The placenta is a fundamental organ for human reproduction, facilitating fetal growth via the exchange of oxygen, nutrients and waste products between mother and fetus, by means of a dense network of fetal villi bathed in maternal blood that flows through the intervillous space (IVS). However, despite its role in adverse pregnancy outcomes associated with impaired maternal-fetal transfer, the influence of placental structure on maternal haemodynamics and the delivery of oxygen and nutrients to the developing fetus is not well understood.

This study employs computational fluid dynamics within physiologically informed 2D and 3D whole-organ-scale placental geometries, whose features are informed by recent *ex vivo* experimental and *µ*CT data, to examine comprehensively the influence of placental anatomy on maternal flow and transport in the IVS. In particular, we consider in detail the impact of the number and placement of maternal decidual arteries and veins that supply and drain the IVS, the location and height of so-called septal walls that loosely separate the placenta into functional units (cotyledons) and sub-units (lobules), and the density of the fetal villous trees as reflected in the rate of uptake of dissolved solutes from the maternal blood and the resistance to flow.

We first exploit the computational efficiency of simulation in a representative 2D geometry to study in detail the sensitivity of haemodynamic markers to these parameters. These results guide our 3D study which reveals that the flow, transport and oxygen uptake are strongly influenced by placental structure, and exposes the vein-to-artery ratio as a key indicator of placental efficiency, regulating a trade-off between a preferential maternal flow environment and fetal oxygen uptake. Conversely, the location and height of the septal walls, a feature that is not well studied, have minimal systematic impact on the macroscopic haemodynamic and transport measures considered here. We also introduce a reduced model, for which analytical progress can be made, and demonstrate its utility in exposing key drivers of maternal-fetal transport.

**Author summary:** In our study, we explored how placental structure affects maternal blood flow and the delivery of oxygen to the fetus. The placenta is a complex and vital organ, but we do not fully understand how its morphology influences its functional efficiency. This is a critical gap in our knowledge as placental blood flow is implicated in serious pregnancy complications such as fetal growth restriction and pre-eclampsia.

To investigate the effect of structure, we used detailed 2D and 3D models of the placenta. These models allowed us to simulate maternal blood flow and oxygen transport therein while changing various structural features, such as the number and location of placental veins and the density of the intervillous space. We found that the ratio of veins to arteries is a key factor in determining the maternal blood flow speed and the amount of oxygen the fetus receives. A higher number of veins relative to arteries helps create the slow-flow environment necessary for a healthy pregnancy, but an excessive ratio can reduce the efficiency of oxygen uptake. Our findings suggest that the vein-to-artery ratio could be an important indicator of placental health, offering a new perspective for understanding and enabling effective early intervention treatments of pregnancy complications.

## Introduction

The placenta is fundamental to the human reproductive system, acting as the interface between the mother and fetus and facilitating exchange between maternal and fetal circulatory systems. The mechanisms governing placental haemodynamics, and their influence on maternal-fetal exchange, are poorly understood, despite inadequate haemodynamics being implicated in multiple maternal and fetal conditions associated with poor pregnancy outcomes, including stillbirth, pre-eclampsia [1] and fetal growth restriction [2]. Furthermore, such poor pregnancy outcomes are correlated to an increase in the risk of pregnancy loss in future pregnancies [3, 4].

The placenta is a complex medium which exhibits various physical processes at different scales, ranging from the macroscopic organ scale [5], through the cell scale [6], to the microscopic molecular scale [7]. A schematic of the placenta, illustrating the key structural features influencing maternal blood flow dynamics and solute transport is provided in Figure 1. Maternal blood flows through the placenta, bathing a dense network of fetal villi across which exchange occurs. On the macroscopic organ scale, cotyledons separate the basal plate into distinct regions. Each cotyledon contains a set of lobules (also known, primarily in the mathematical literature, as placentones) where each lobule may contain a chorionic stem villus and/or spiral artery. Both cotyledons and lobules are delineated by septa on the basal plate protruding within the placenta. High-speed maternal blood is brought into the placenta through the decidual spiral arteries that enter through the basal plate. This blood travels into a small free-flow region, devoid of fetal villi, known as the central cavity (CC) [8]. Once there, it undergoes a rapid transition from a fast inertia-dominated flow to a slow viscous-dominated flow as it saturates the low-permeability porous medium surrounding the CC. This materno-placental medium is formed by the dense fetal villous tissue and is known as the intervillous space (IVS). Exchange of nutrients and oxygen from maternal to fetal blood primarily takes place via diffusion across the walls of terminal villi, small and peripheral villous branches of the villous tree containing fetal capillaries, and is subsequently transported back through the tree and eventually to the fetus via the umbilical cord; waste products are transported in the opposite direction. Here, we consider the placental structure to encompass both the macroscopic anatomy (cotyledon, lobule and septum layout) and the maternal vasculature (arterial inflow, venous return).

**Fig 1.**
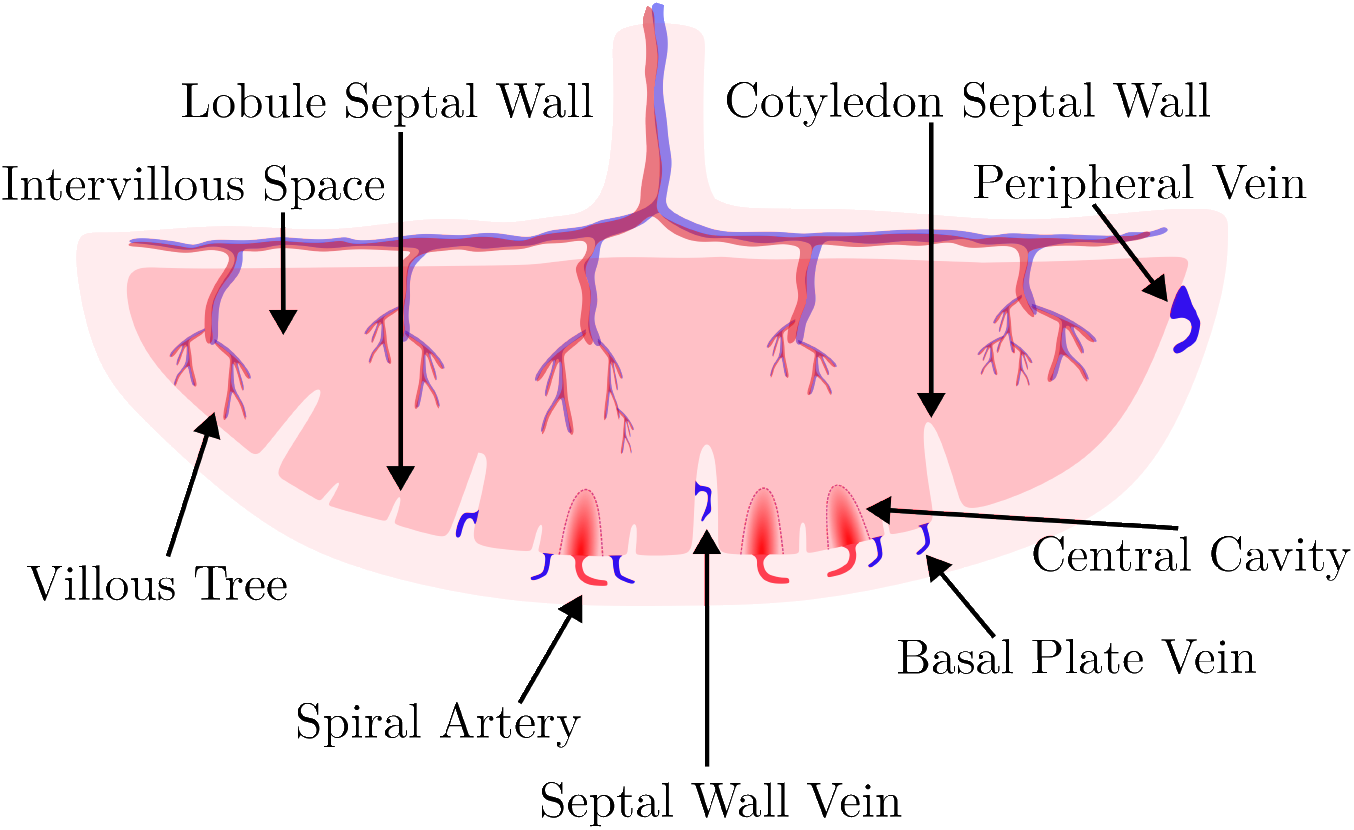
Schematic of a 2D cross-section of a placenta. Maternal blood enters the central cavities from the spiral arteries where it rapidly decelerates as it enters the intervillous space. In this viscous-dominated flow regime, oxygen and nutrients are transferred from maternal blood to the fetal vasculature before exiting the placenta via venous return. Five cotyledons are shown here, each with varying numbers of lobules, arteries, veins.

For healthy placentas, the flow experiences high resistance with large pressure drops in transitioning from the spiral arteries to the IVS [9] [10, p 33]. Once inside the IVS, the flow is characterised by a preferential ‘slow flow’ regime which balances potentially damaging viscous energy expenditure (and associated shear stresses exerted on the villi) in the dense villous IVS with exchange of nutrients and oxygen to the fetal vasculature. Additionally, it has been observed that regions of low oxygen concentration gradient have higher densities of terminal villi, with lower blood flow speeds believed to contribute to their growth due to increased exposure time resulting in greater nutrient and oxygen exchange [11]. Conversely, high flow speeds are believed to be responsible for reduced oxygen uptake in the placenta due to increased damage to the villi and reduced oxygen exchange between the maternal blood and villi; consequently the maternal blood leaves the placenta with a higher-than-normal nutrient and oxygen concentration [12].

In compromised placentas, characteristics of this optimal flow regime may be absent [5, 13, 14]. Hence, flow within the placenta and IVS which exhibits dynamics outside of the slow flow regime may suggest inadequate functioning of the placenta. For example, in pre-eclampsia incomplete spiral artery remodelling leads to increased flow speeds and thereby greater inertial effects within the IVS [12]. Correspondingly, higher viscous dissipation is likely to be observed in the IVS in comparison to healthy placentas. Higher shear stresses have been found to inhibit cellular activity responsible for the remodelling of the spiral arteries in the first trimester [15]. Thus, the rapid transition from fast CC flow to a slow viscous-dominated flow in the IVS is crucial in creating a low energy expenditure environment, characterised via low viscous dissipation within the IVS which reduces shear stress on the villi while enabling nutrient and oxygen exchange.

One of the primary challenges in studying placental haemodynamics is the complex and variable nature of its structure, with significant morphological and physiological changes occurring through gestation. *In vivo* investigation is severely hampered by self-evident safety and ethical considerations and the limited utility of (non-primate) animal models. It is unique, however, in offering convenient *ex vivo* experimentation after delivery so that physiological examination, and transport and flow experiments can be undertaken on a living human organ, albeit in the presence of some unavoidable structural changes [16]. Non-invasive techniques such as MRI offer a powerful route to probe placental structure, flow and transport, and moreover is suitable for repeated scanning, thereby allowing investigation of placental function longitudinally across gestation [17, 18]. However, inferring the detail of IVS flow patterns and oxygen transport from standard MRI remains an open challenge; see [19–21], for example, for some recent studies on placental MRI. Confounding factors include disambiguating maternal and fetal flow, sub-MRI voxel-scale complexity, and the potential non-uniqueness of the inverse problem associated with directly determining such flows from data.

Mathematical and computational modelling provides a natural route to understanding placental haemodynamics, allowing comprehensive study of flow, transport and exchange at levels inaccessible to experimental investigation. From a fluid-mechanical and multiphysics standpoint, the variable and multiscale nature of the placenta offers a diverse range of problems in IVS flow, from delivery to (and transport across) terminal villi at the microscale to macroscale IVS flow in porous media, and a host of problems in between. Our focus is on modelling macroscopic IVS flow and transport and so we do not seek to provide a review of these problems here, nor of those pertaining to fetal flow; the interested reader is directed to [22] for detailed coverage.

The first computational study of IVS flow was provided by [23], in which a single lobule was idealised as a square domain, with maternal flow governed by Darcy’s law and a spatially-varying permeability field accounting for the low-permeability CC region. This Darcy flow approach was adopted more recently by [24] to obtain analytical expressions for the blood flow field in a 3D hemispherical domain to study how vein placement and CC size affects flow and transport, exposing in particular potential ‘short-circuiting’ behaviour between arterial supply and venous drainage in this model. Philosophically similar studies have been presented by [25] in 2D, with a focus on wall shear stress (together with other more microscale studies) and by [26] in 3D in which the influence of arterial jets was considered.

A common theme of the above-noted and other studies is the adoption of geometrically-simplified representations, typically of a single lobule, for analytic tractability or computational expediency, thereby prohibiting a clear understanding of the influence of its complex structure on flow and transport phenomena and how this translates to the whole-placenta scenario. These considerations are especially important given the structural complexity of the placenta, its variation across different pathologies, such as pre-eclampsia, fetal growth restriction, and diabetes, and significant inter-subject differences. Moreover, the detail of placental structure is not well-characterised in general. By way of example, estimates of spiral artery number range from 30 to 150 [12, 24, 27] and information on the number of veins is scarcer still, with [24] giving a range of 20 to 200. In each case, placement (as highlighted by [24]) will strongly influence flow dynamics; this, and the significance of septal wall veins in IVS flow, are not yet studied.

Here, we seek to address this deficiency, providing a comprehensive computational study of the influence of placental structure on IVS flow and solute transport within physiologically relevant geometries that exploit recent *ex vivo* physiology investigations and *µ*CT data [28]. A systematic sensitivity study on the parameters which characterise placental geometries, and flow and uptake model characteristics, allows us to probe their individual and collective impacts on placental haemodynamics and solute delivery. Through this approach, we aim to provide new understanding of the critical determinants of blood flow dynamics and oxygen transport within the placenta.

## Methods

In this section, we outline the models which describe the placental flow problem. From here on, dimensional variables are denoted with asterisks and dimensionless variables without. Specification of the characteristic values we use for the non-dimensionalisation can be found in **S1 Parameters and Geometry Description**.

### Ethics Statement

Women with singleton pregnancies were recruited after informed written consent at the Queen’s Medical Centre, Nottingham University Hospitals NHS Trust, UK, for the SWIRL (Stillbirth, When Is Risk Low?) study, with IRAS project ID 321374 / REC (23/EM/0052) under the overarching Wellcome-Leap In Utero project. Recruitment was carried out in accordance with The Code of Ethics of the World Medical Association (Declaration of Helsinki). The representative term placentas shown in **S1 Parameters and Geometry Description** were obtained following elective Caesarean section (≥ 37 weeks’ gestation). Placentas were collected immediately after delivery with the cord clamped. Participants were of similar age, non-smokers, and had uncomplicated pregnancies.

The *µ*CT data described in **S3 Homogenisation** was acquired under ethical approval from the ethics committee at the University of Manchester (Human Research Authority Reference 24/NW/0104 and DAPHNE, REC 22/YH/0144).

### Placental Geometry

Here we describe the model geometry that we employ in our 3D computational study, which is specified to accommodate the key structural features described above. Figure 2 illustrates the pipeline used for generating our computational geometries, which can be summarised as follows.

**Fig 2.**
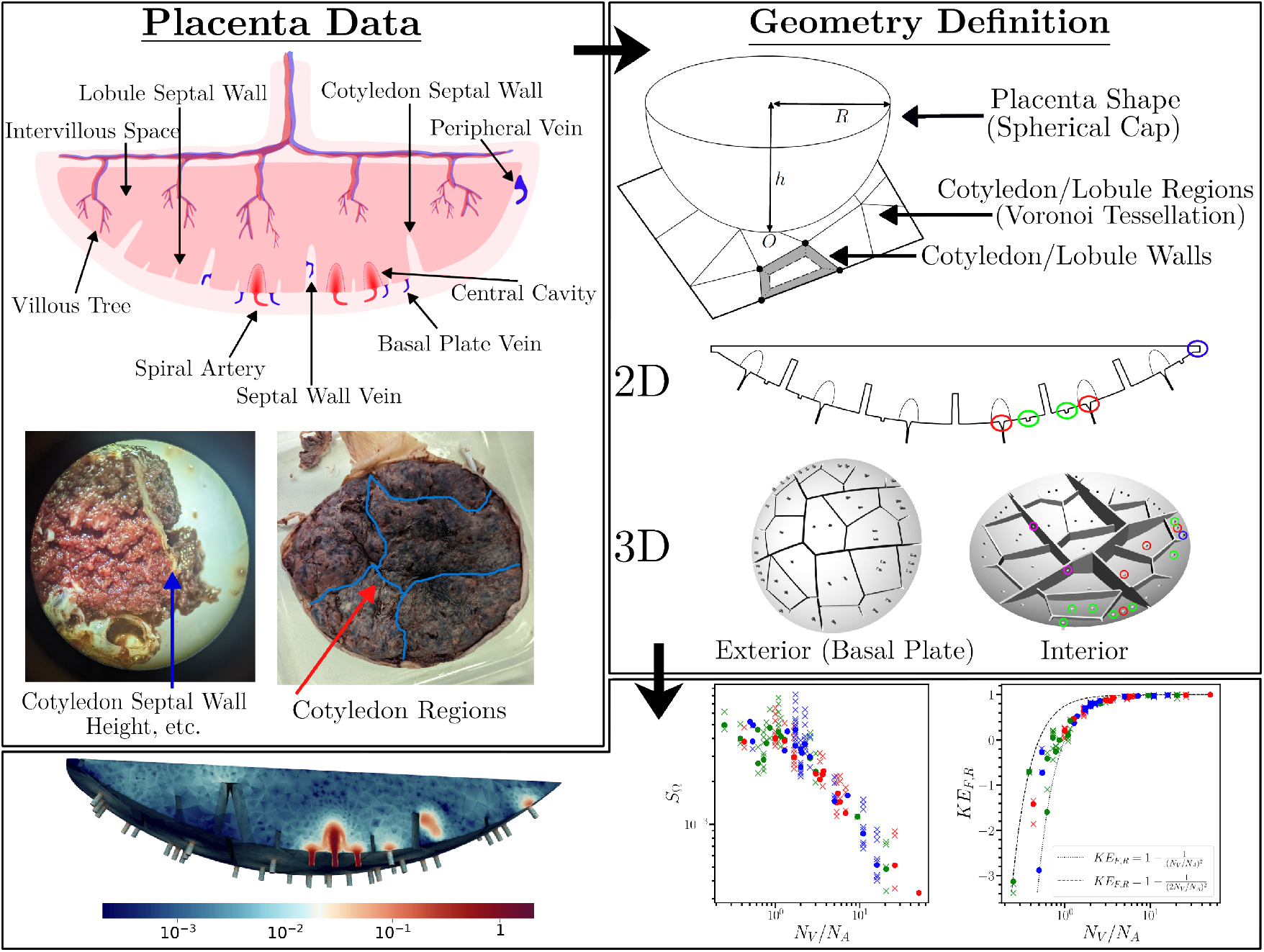
Illustration of the geometry creation process. Step 1: data from placental measurements are used to define key features of the placental geometries. Step 2: the placental cotyledon geometry is generated by a 2D Voronoi tessellation which is projected up through the spherical cap, hence creating the septal wall structures. The black dots are the vertices of a convex Voronoi cell which forms either a cotyledon or a lobule. The grey area between cells denotes the septal wall. The final reconstructed geometries in 2D and 3D are shown with examples of different types of vessel highlighted: spiral arteries (red circles), basal plate veins (green circles), peripheral veins (blue circles) and septal wall veins (purple circles). Step 3: A cross-section of the 3D flow solution, alongside exemplar results (see Section **Results**) showing how the performance markers respond to changes in the geometrical parameter space.

- The placenta is represented as a spherical cap, characterised by base radius *R* and volume 𝒱 = *πh*^2^(*R* − *h/*3), in which *h* denotes placental height. The curved surface and flat parts of the resulting geometry represent the basal and chorionic plates of the placenta, respectively.
- Lobe regions are constructed using placental measurements taken from *ex vivo* data: septa, delineating cotyledon boundaries, are marked on images of delivered placentas, which are used as the basis for a Voronoi tessellation [29], whose projection into the 3D geometry defines cotyledon septa of height *H*_*C*_ and thickness *T*_*C*_; lobules, with wall height *H*_*L*_ and thickness *T*_*L*_ are defined via a similar process, but their positions within lobules are selected randomly since their boundaries are less clearly defined on the surface of the *ex vivo* placenta.
- The maternal vasculature comprises *N*_*A*_ decidual spiral arteries, which supply maternal blood to the materno-placental domain. This blood first enters the CC, which is modeled as a semi-ellipsoidal region with lengths of the minor and major axes *a, b*, respectively, centered on the entry point of each spiral artery. Blood then traverses the IVS (defined here to be the remaining part of the interior of the placenta), exiting through *N*_*V*_ = *N*_*S*_ + *N*_*B*_ + *N*_*P*_ decidual veins, where *N*_*S*_, *N*_*B*_ and *N*_*P*_ denote the number of veins located in septal, basal plate, and peripheral locations, respectively. The numbers and positions of decidual arteries and veins are selected at random.

For a given *ex vivo* placenta, specifying the geometrical parameters noted above thereby provides a realisation of a representative placental geometry in which we study maternal flow and associated solute transport.

We remark that in the numerical explorations that follow, we first exploit a representative 2D geometry (generated in an analogous manner; see [30]) for computational efficiency, allowing us comprehensively to explore the geometrical and model-parameter space, prior to the focused interrogation of our 3D placental domain. Further information on the 2D and 3D geometrical setups, including values for the aforementioned features, is given in the Supporting Information **S1 Parameters and Geometry Description**.

### Mathematical Model of Flow, Transport and Uptake

We assume that the flow of maternal blood is isothermal with the fluid described as incompressible and Newtonian, which is governed by Navier–Stokes–Darcy equations

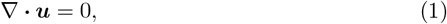

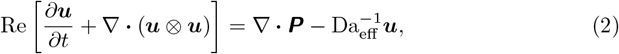

where ***u***(***x***, *t*) denotes the maternal blood velocity at position ***x*** = (*x, y, z*) and time *t*. The fluid stress tensor is denoted ***P*** = ∇***u*** + (∇***u***)^*T*^ − *p****I***, with pressure *p* and identity tensor ***I***. Flow characteristics are determined by the dimensionless Reynolds (Re) and effective Darcy (Da_eff_) numbers. The distinction between the CC and IVS flow is characterised by a spatially varying effective Darcy permeability coefficient. We model the transition as a finite-thickness layer between the CC, where the Darcy term is negligible, and the IVS, where the Darcy term dominates (in which the value of Da_eff_ is estimated from *µ*CT data by means of asymptotic homogenisation, as described in Section **IVS Permeability**, below). This provides a convenient way to describe the increase in density of villous architecture, while avoiding the introduction of sharp interfaces between free flow and porous flow regions that are commonly employed in existing models [26, 31]. It also avoids the requirement to define and approximate appropriate interface conditions, which is advantageous from the point of view of the numerical methods that we subsequently employ.

Maternal blood transports a passive diffusible solute, governed by an advection-diffusion-reaction equation,

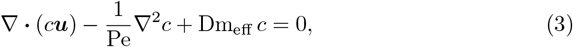

where *c*(***x***, *t*) denotes the oxygen concentration. Solute transport characteristics are determined by the dimensionless Péclet (Pe) and effective Damköhler (Dm_eff_) numbers, with the latter being spatially varying, in the manner of the Darcy number above, to distinguish uptake dynamics in the CC from those in the IVS. We note that due to the relatively low pulsatility index of the third trimester pregnancies [13], we consider the time-independent oxygen transport problem. This model then provides a simple and numerically tractable representation of the delivery of oxygen or nutrients from the maternal supply to the developing fetus via the placental villous tree. For brevity, we henceforth refer to *c* as the oxygen concentration, while recognising the complexity associated with more complete descriptions of oxygen uptake and fetal-maternal transport. Consistent with our representation of villous density, uptake by the villi is modelled as spatially varying linear reaction, as explained in more detail in Section **IVS Permeability** below.

The maternal flow domain is denoted by Ω and the *N*_*A*_ artery inlet surfaces by Γ_in,*i*_, *i* = 1, …, *N*_*A*_ and are treated as inflow boundaries. We denote the full inflow boundary by Γ_in_ with 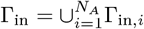. Similarly, the *N*_*V*_ vein surface faces are treated as outlets with the outflow boundary 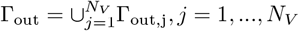. The remaining surface is denoted by Γ_D_; the inward-pointing unit normal to *∂*Ω is denoted by ***n***.

Suitable boundary data are given by

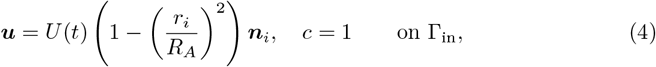

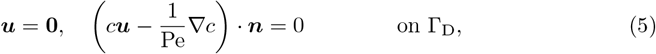

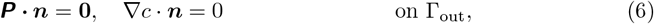

in which *r* is the local radial coordinate in the plane perpendicular to the inward-pointing unit normal ***n*** and centred on the origin of the circle defining each arterial inlet Γ_in,*i*_. The inlet (and outlet) faces are assumed to be flat, so the relevant normal vectors are constant. In 2D, we assume the flow is steady with equivalent time-independent boundary conditions applied.

These conditions describe the entry of maternal blood, carrying solute at dimensionless concentration *c* = 1, to the placenta through arteries of radius *R*_*A*_ via a pulsatile Poiseuille inlet flow, whose amplitude *U* (*t*) is obtained from Doppler ultrasound measurements [32] (scaled such that the systolic peak is as measured in third trimester [33, 34]). The pulsatile inlet wave form which provides *U* (*t*) is shown in Figure 3. On the placental walls, no-slip and no-penetration are enforced, while a free-flow condition is applied at venous outlets.

**Fig 3.**
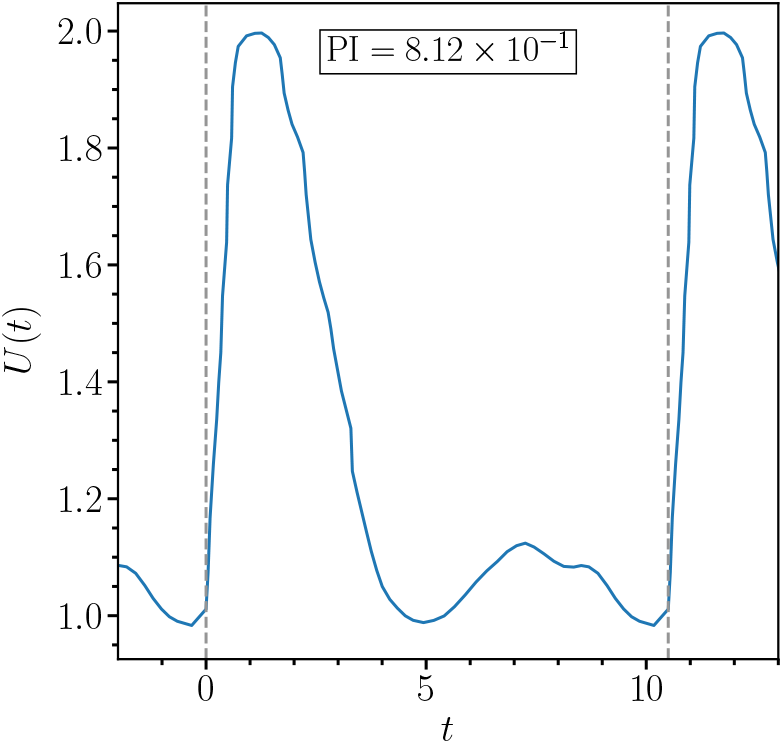
Scaled pulsatile waveform that specifies the peak velocity through every spiral artery. The grey vertical lines indicate the beginning and end of a cardiac cycle with the pulsatility index (PI) of the waveform shown in the legend.

We take the initial condition as ***u*** = **0**. Due to the viscous-dominated regime of the flow in the IVS, the flow rapidly converges to periodicity with respect to the cardiac cycle, see **S2 Cardiac Cycle Convergence** for further information. Hence, we are able to use the flow field generated during the second cardiac cycle for our transport and uptake computations. In particular, when calculating the performance markers we consider the time-averaged mean over the second cardiac cycle. For the steady state transport problem, we use the flow field corresponding to the peak inlet velocity from the second cardiac cycle. The advantage of this approach is to enable a reduction of the time-dependent quantities into single representative ones.

The nondimensionalisation, and the geometrical and other model parameters, together with their sources, can be found in **S1 Parameters and Geometry Description**. We emphasise that in the numerical explorations that follow, we vary these parameters across a wide range to explore in detail those features controlling system behaviour; physiologically reasonable estimates are indicated in **S1 Parameters and Geometry Description** and referred to in the main text as ‘nominal’ values.

### IVS Permeability

Estimates for the permeability of the IVS, a key determinant of the flow and transport dynamics, vary in the literature. To obtain a suitable value for our study, we employ *µ*CT scans of an *ex vivo* placenta and a classical two-scale asymptotic homogenisation process, as described in [35, 36] and elsewhere (though we emphasise that we vary the permeability, and other parameter values, as part of our study). Detail is given in the Supplemental Information **S3 Homogenisation**, but we describe briefly the method here for completeness.

We consider the IVS as an idealised porous structure, represented as a highly connected material with spatially periodic microstructure, saturated with a viscous Newtonian fluid, and characterised by two asymptotically well-separated lengthscales. This scale separation allows a leading-order tissue-scale Darcy permeability to be obtained from a Stokes problem posed on a periodic domain associated with the pore-scale flow domain surrounding terminal villi.

We exploit *µ*CT data of an *ex vivo* placenta to construct our microscale flow domains. See **S3 Homogenisation** for details of the data acquisition; in brief this data comprises 2120, 2520 and 2520 voxels in the *i, j, k* coordinate directions, respectively, with each voxel representing either IVS flow domain or villi. As there is no maternal blood flow within the villous domain, we take this as impermeable solid. We sample a domain *Ŷ* = [0, 1*/*2]^3^ from this data using the same number of voxels in each coordinate axis. The relevant flow domain is denoted by *Ŷ*_*f*_ ⊂*Ŷ*. Reflection of *Ŷ*_*f*_ against each coordinate axis provides a periodic unit cell *Y*_*f*_ ⊆ [0, 1]^3^. Solution of a standard forced periodic Stokes problem on this domain provides a local flow; integration over the unit cell to remove local variation yields the macroscopic permeability tensor.

The classical homogenisation method we employ here does not account for the statistical variability in the underlying geometry that may be of importance in accurately characterising the flow in the IVS (see, for example, [37] for detailed consideration of placental transport and uptake in the presence of spatial disorder). Nevertheless, it provides empirical support for our estimate, thereby locating this parameter in a physiologically reasonable region of the parameter space. To obtain a physiological estimate for the IVS permeability tensor, we compute this quantity from a range of domains of differing sizes sampled from the *µ*CT data to obtain convergence of the predicted value.

For simplicity, in the simulations that follow we reduce the permeability tensor, via the Euclidean norm, to a scalar value *κ*^*^, with 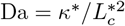 for characteristic length scale 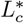 (see **S1 Parameters and Geometry Description**), to use as the value for IVS permeability. Figure 4 provides an illustration of the data, together with examples of these computed values on the finest data sample we used. In **S3 Fig 1**, additional plots are shown for increasingly many voxels, showing a convergent trend for increasing voxel number. Guided by this, we take the nominal value as Da = 10^−4^ (*κ*^*^ = 10^−8^ m^2^).

**Fig 4.**
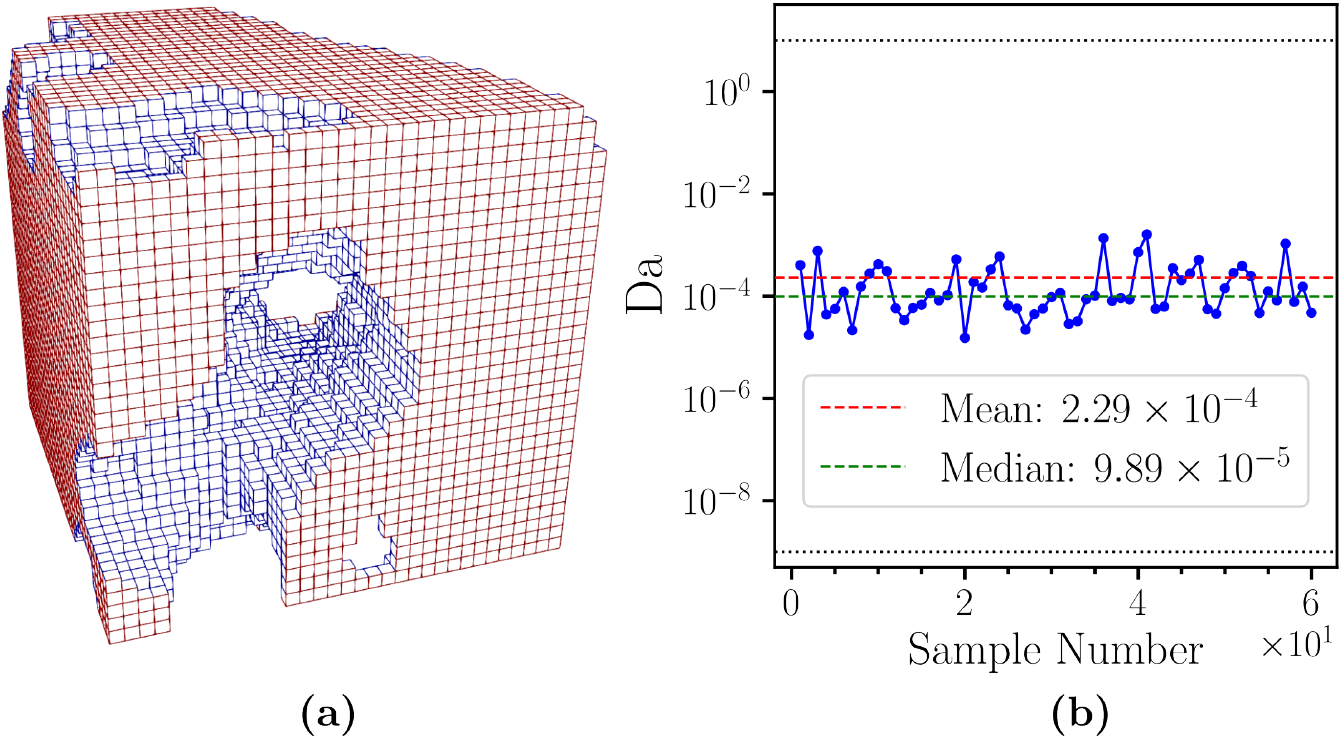
(a): An example of a unit cube geometry where hollow regions represent villous structures and cubes represent the flow domain, (b) Indicative results of IVS permeability (Da) computed from a range of unit cubes of size 512^3^ voxels (‘sample number’), sampled from *µ*CT data [28].

We reiterate that to model the distinction between CC and IVS regions, we employ a spatially varying IVS permeability (and associated uptake rate) such that

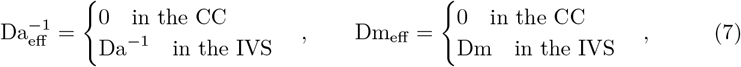

with the transition between these values smoothed over a finite region by means of a smoothing function:

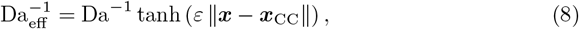

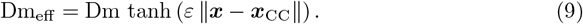

For a given ***x*** in the flow domain, ***x***_CC_ is the point that minimises the distance from ***x*** to any CC boundary and *ε* is the smoothing coefficient which determines the thickness of the transition region. We take *ε* = 3.68, which ensures the transition region is suitably thin and resolved on our computational meshes.

### Coupling to the Feto-placental Circulation

In our model of maternal blood flow and solute transport, solute exchange between the maternal and fetal blood is included through an uptake term, Dm_eff_ *c*, in (3). Further insight can be gained by applying a homogenisation argument to the solute transport and exchange dynamics from the perspective of the feto-placental circulation. By extending this analysis to the maternal transport-exchange system, we can derive a reduced model of solute uptake which we also use to validate the detailed placental flow model.

To this end, we assume that fetal placental tissue forming the solid phase of the porous medium is predominantly formed from terminal villi. A villus occupying a representative elementary volume (REV) at ***x***^*^ of volume 𝒱^*^ (where 𝒱^*^ is comparable to *κ*^*3*/*2^ in magnitude; see **S1 Parameters and Geometry Description**) is supplied by fetal circulation that delivers solute to the villus at a (fetal arterial) concentration 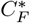 with a local volume flux 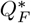 of fetal blood through the capillary network of the villus, returning blood with concentration 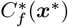 to the venous side of the feto-placental circulation. We assume that villi operate independently of their neighbours and that an average villus can be characterised by a lengthscale ℒ^*^, which measures diffusive exchange capacity, and a tissue diffusivity 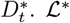 can be interpreted as a ratio of surface area for exchange within the REV to diffusion distance between maternal and fetal blood [38]; it can be expected to scale like *κ*^*^*/h*^*^ for some diffusion distance 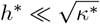. Transport within the villus is then characterised by a fetal Damköhler number 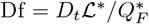, which distinguishes fetal-exchange-limited transport (Df ≪ 1) from fetal-flow-limited transport (Df ≫ 1). Note that, although the regime Df ≪ 1 is constrained by a diffusive process across the villous tissue separating maternal and fetal blood, it is referred to here as fetal-exchange-limited, both for brevity and to avoid confusion with the diffusive processes modelled within the maternal blood in (3).

The time-averaged maternal concentration field *C*^*^(***x***^*^) is assumed to vary smoothly over multiple REVs, satisfying ∇ ^*^· (***u***^*^*C*^*^ − *D* ^*^*C*^*^) = −*N* ^*^. Following [38], the (homogenised) flux of solute per unit volume between maternal and fetal blood can be written

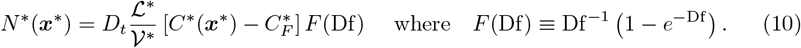

The function *F* approximates the transition between exchange-limited and fetal-flow-limited transport, i.e. *F* decreases monotonically from *F* ≈ 1 for Df ≪ 1 to *F* ≈ Df^−1^ for Df ≫ 1. The corresponding fetal solute flux emerging from an individual villus is 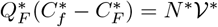. Thus the fetal venous concentration and the maternal IVS concentration are related by

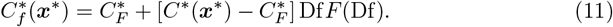

Setting 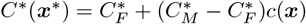, where 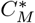 is the maternal arterial concentration, we recover (3) by taking the uptake rate to be

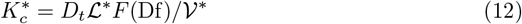

in the definition 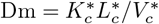 in **S1 Table 1**. The characteristic velocity 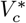 in **S1 Table 1** is taken to be the half-maximal inlet flow speed; the arterial flow speed, averaged over a period *T* and over an arterial cross-section, is 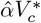, where 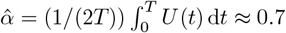, so that the total time-averaged maternal arterial volume flux is 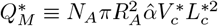, where *R*_*A*_ denotes the (dimensionless) arterial radius. Using (10) and the definition of *C*^*^(***x***^*^), the rate at which solute is delivered from mother to fetus is then

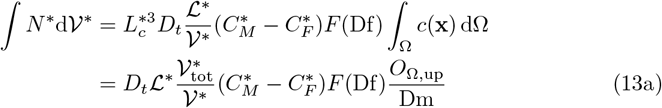

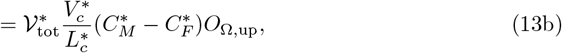

where the factor of 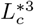 arises from the dimensional integration domain 𝒱^*^ and 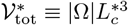 is the total volume of the domain occupied by fetal tissue. In (13a), *O*_Ω,up_ ≡ |Ω| ^−1^ ∫_Ω_ Dm *c* dΩ measures global solute uptake, one of the key flow and transport performance markers we investigate in our computational studies (33), and we recognise 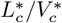 in (13b) as a characteristic timescale (**S1 Table 1**).

The ratio 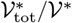 in (13a) is a measure of the number of exchange units (terminal villi) within the placenta and ℒ^*^ is a measure of the exchange capacity of each unit. Thus *F* (Df)*O*_Ω,up_*/*Dm in (13a) measures the proportion of the placenta’s total exchange capacity 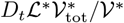 that is recruited for exchange. Under conditions of fetal-exchange limitation, for which *F* = 1, *O*_Ω,up_*/*Dm ≤ 1 measures the degree of maternal flow limitation. The dependence of *O*_Ω,up_ on Dm is evaluated computationally in **Results** below; this relationship implicitly encompasses dependence on fetal transport via the relation

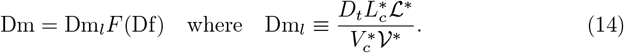

Here, Dm_*l*_ is a reference maternal Damköhler number that is independent of fetal conditions; the subscript *l* recognises its relevance to local (villus-scale) exchange.

Equation (13b) also provides an interpretation of *O*_Ω,up_ under conditions of maternal flow limitation. Suppose again that *F* = 1, *i*.*e*. from the perspective of the fetus the uptake is exchange-limited, but that the maternal flow is so weak that the full inlet flux is taken up by the fetal circulation. In this limit

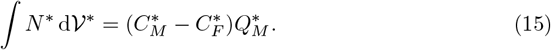

Thus for sufficiently large Dm, using (13b) *O*_Ω,up_ asymptotes to

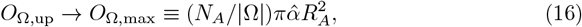

a ratio connecting the number of arterial sources *N*_*A*_ to the approximate number of lobules 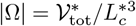.

### Reduction of the Exchange Model

A preliminary estimate of the transition between maternal flow limitation and maternal exchange limitation is provided by comparing (15) to the total exchange capacity (13a), i.e. 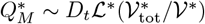, or equivalently

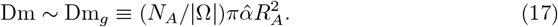

This describes the case for which the (local) maternal Damköhler number Dm = Dm_*l*_*F* (Df) reaches a threshold determined by the (global) ratio between the number of arteries and the approximate number of lobules 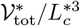.

A candidate approximation for the mean solute uptake is provided by

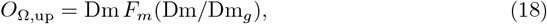

with *F*_*m*_ given, for example, by *F* in (10). This has the anticipated limits *O*_Ω,up_ ≈ Dm for Dm ≪ Dm_*g*_ (maternal exchange limitation) and *O*_Ω,up_ ≫ *O*_Ω,max_ for Dm Dm_*g*_ (maternal flow limitation). The approximation (18) neglects additional dependence on other parameters, such as the ratio *N*_*A*_*/N*_*V*_, so we fit a more general ansatz (which reduces to (10) with specific parameter choices) when comparing with our simulation results on complex geometries.

The analysis above allows us to combine maternal and fetal perspectives: when measured with respect to the overall exchange capacity of the placenta, net solute exchange can be estimated in terms of the independent parameters Df and Dm_*l*_*/*Dm_*g*_,

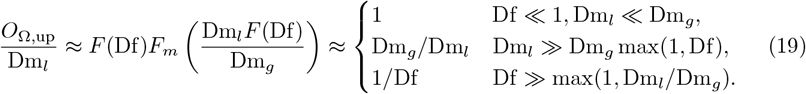

The three limits define respectively exchange-limited transport, maternal-flow-limited transport and fetal-flow-limited transport. A transition between maternal- and fetal-flow-limitation is anticipated when

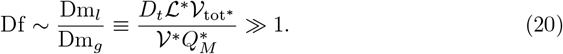

### Numerical Method

For the spatial discretisation of (1–3), a symmetric interior-penalty discontinuous Galerkin (DG) method is used [39, 40]. In 3D, a residual-based artificial viscosity term is included in the transport problem [41] to damp numerical oscillations, with the viscosity parameter taking the modified form proposed in [42]. We use simplex meshes and employ piecewise discontinuous linear polynomials to approximate the velocity and concentration, and piecewise constant polynomials to discretise the pressure. For the time discretisation of (2), we use a semi-implicit BDF2 scheme, wherein the nonlinear transport term is linearised via a Newton–Gregory extrapolant [43]. In 2D, as we only consider steady flow, Newton’s method is employed to compute the solution of the underlying system of nonlinear equations. The flow solver employed in this article has previously been developed and validated in, for example, [44].

### Flow and Transport Performance Markers

We interrogate placental flow and transport dynamics under variation of the geometrical and model parameters by means of a range of macroscopic haemodynamic performance markers, these being chosen as indicative of placental (dys)function. We place particular emphasis on measuring the effectiveness of the IVS in creating the slow flow regime (via placental speed and energy markers) and the flow and exchange process within the IVS (via viscous dissipation and oxygen uptake). For time-dependent quantities, we additionally time-average over a cardiac cycle^11^.

The energy-related markers we consider below are derived from (1) and (2). The relevant energy equation is

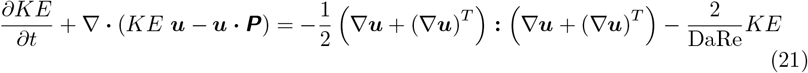

where *KE* denotes the kinetic energy density,

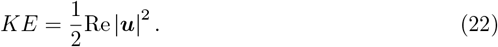

#### Mean blood flow speed, *S*_Ω_, is defined by

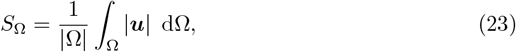

and quantifies how effectively the placenta is able to slow the flow down in the volume Ω via resistivity of the medium to the flow and potentially reducing damaging shear forces in Ω.

**Cross-cotyledon flux**, 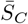, is defined by

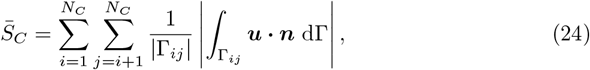

where Γ_*ij*_ = Ω_*i*_ ∩ Ω_*j*_ represents the interface between cotyledons *i* and *j*. It measures the overall cross-cotyledon flow of maternal blood and indicates how well the maternal blood is able to perfuse through the IVS, and the extent to which each cotyledon acts as an independent exchange unit.

**Slow flow volume ratio**, 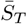, is defined by

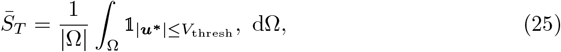

where 𝟙 is the indicator function, ***u***^*^ is the dimensional velocity and the slow flow velocity threshold value has been chosen from [5] to be *V*_thresh_ = 5 *×* 10^−4^ m s^−1^. This measures the percentage of the placental volume where slow flow is present, so a larger value of 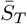 corresponds to a greater proportion of the placenta experiencing a preferential flow regime.

**Maximum mean arterial pressure drop**, Δ*p*_*M*_, is defined by

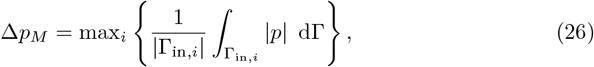

and measures the pressure drop within the placenta, and hence the force of the fluid in the CC required to overcome the resistance of the IVS.

**Kinetic energy**, *KE*_*F,R*_, and **total energy**, *E*_*F,R*_, **flux loss ratios** are defined by

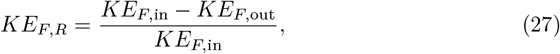

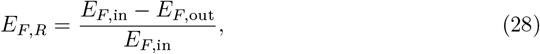

where

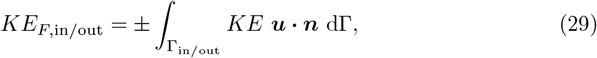

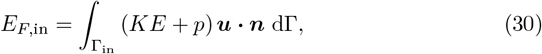

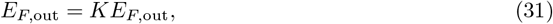

and ***n*** is an inward-pointing unit normal vector. These measure the efficiency of the placenta in removing energy from the flow through viscous stresses to create a preferential slow flow environment. For *KE*_*F,R*_, an approximation for its dependence on the numbers of veins *N*_*V*_ and arteries *N*_*A*_ can be obtained by appealing to conservation of mass and the assumption of Poiseuille in- and out-flow through arteries and veins of uniform radii *R*_*A*_ and *R*_*V*_, respectively. This is given by

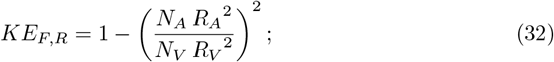

the full derivation is provided in **S5 Kinetic Energy Flux Loss Ratio Approximate Law**. We note that boundary conditions (5) and (6) have been used to derive the simplified forms (30,31).

**Mean oxygen uptake**, *O*_Ω,up_, as defined by

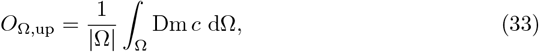

characterises the efficiency of the placenta in delivering oxygen from the maternal blood in the placenta to the fetal vasculature over the volume Ω (see (13)). The reduced exchange model (18) provides an analytical estimate of the dependence of *O*_Ω,up_ on Dm which we compare below to our simulation results.

**Viscous dissipation rate in the IVS**, 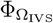, is defined from (21) by

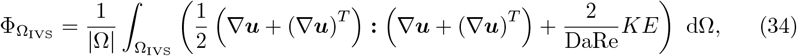

which measures the damaging shear forces exerted on the villi in the IVS due to viscous effects. The second term on the right hand side, known as the Darcy dissipation rate, dominates the contribution to 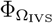 for physiological Da.

## Results

### Preliminary 2D Simulation Study

Our 3D computational study has been guided by a comprehensive set of simulations in 2D [30], whose relative computational tractability allows comprehensive coverage of the parameter space. Overall, in 2D we have data from 9400 simulations, a far higher number than is achievable in 3D.

In figures 5 and 6 we summarise the key findings from the 2D parameter study. In both sets of plots, the dashed black line denotes the median. For data exhibiting considerable variability, we additionally plotted the 25^th^ and 75^th^ percentiles and interquartile range shown by the red line, green line and blue area, respectively. Cross markers indicate data points which lie outside the interquartile range.

**Fig 5.**
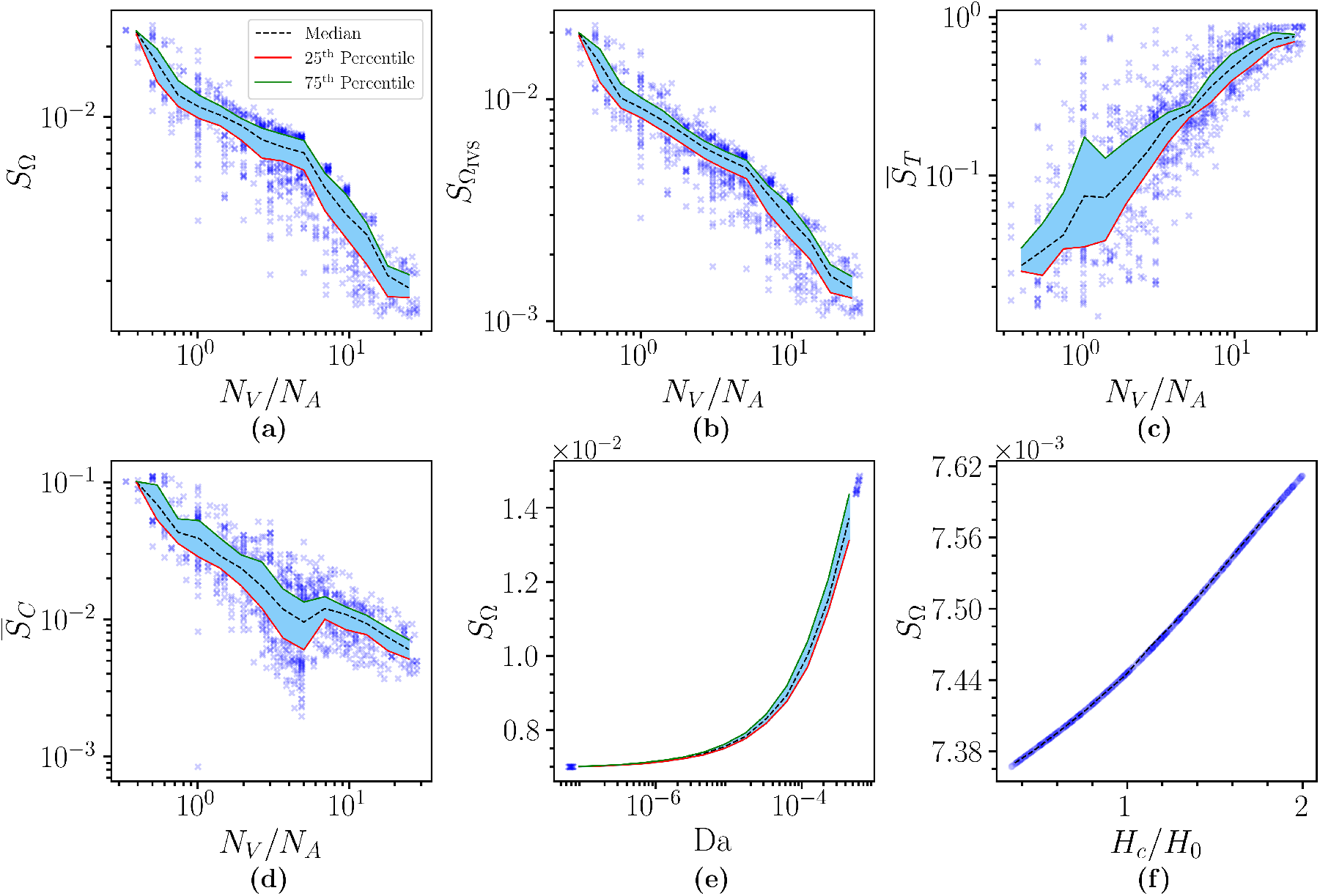
Flow markers in 2D. Plots of selected blood flow metrics against problem parameters in 2D. The impact of the ratio of the numbers of arteries and veins *N*_*V*_ */N*_*A*_ on: (a) the mean flow speed in the total flow domain *S*_Ω_, and (b) the IVS in particular *S*_Ω_, (c) the slow flow volume ratio 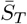 and (d) the cross-cotyledon flux 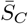. The variation of *S*_Ω_ with (e) Darcy number Da and (f) cotyledon septal wall height, expressed as a relative increase from the nominal heights of the smaller and larger septal walls, *H*_0_ (6.9 mm and 14.07 mm, respectively). Parameters are as given in **S1 Parameters and Geometry Description**, apart from as indicated. In (e,f), the outlet numbers (but not positions) are fixed such that there are no septal wall veins, and one artery and two basal plate veins per cotyledon, for consistency with other studies. The red and green lines represent the 25^th^ and 75^th^ percentiles, respectively, with the blue region between them representing the interquartile range. The cross markers represent data points outside the interquartile range and the black dashed line represents the median of all data points.

**Fig 6.**
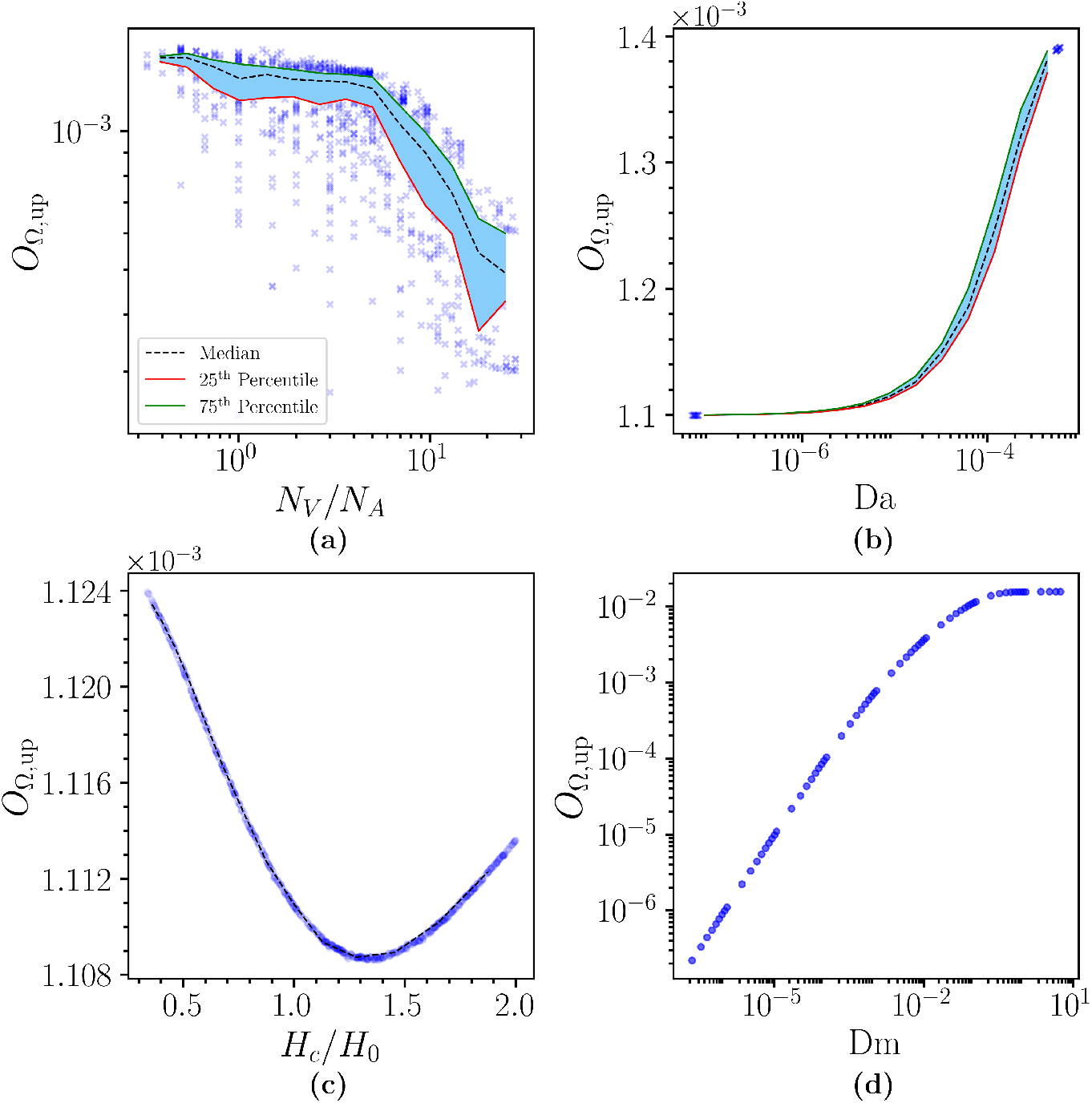
Uptake markers in 2D. Plots of oxygen uptake against (a) the ratio of the number of veins and arteries *N*_*V*_ */N*_*A*_, (b) Darcy number Da, (c) cotyledon septal wall height, expressed relative to the nominal heights of the smaller and larger septal walls, *H*_0_ (6.9 mm and 14.07 mm, respectively) and (d) Damköhler number Dm. In (b–d), the outlet numbers (but not positions) are fixed such that there are no septal wall veins, and one artery and two basal plate veins per cotyledon, for consistency with other studies. The red and green lines represent the 25^th^ and 75^th^ percentiles, respectively, with the blue region between them representing the interquartile range. The cross markers represent data points outside the interquartile range, the circle markers represent all data points and the black dashed line represents the median of all data points. Parameters are as given in **S1 Parameters and Geometry Description**, apart from as indicated.

As expected, we observe that blood flow speed metrics are significantly impacted by *N*_*V*_ */N*_*A*_ (Figure 5(a–d)). An increase in the number of veins or a decrease in the number of arteries results in reduced speed (globally, and within the IVS specifically). As a result, larger regions of the placental domain experience the slow flow regime.

Interestingly, an increase in *N*_*V*_ */N*_*A*_ results in a decrease of cross-cotyledon transport, corresponding to a diminished placental perfusion efficiency, see Figure 5(d), potentially due to lower inertia of the flow and a greater tendency for blood to return via the local venous pathways. We further highlight that for a given value of *N*_*V*_ */N*_*A*_, the flow characteristics vary quite substantially, depending on the artery and vein placement in a given realisation, for example.

Decreased villous density (that is, an increase in Da) corresponds to greater mean placental speeds due to reduced flow resistance in the IVS (Figure 5(e)). Increasing the cotyledon septa height results in increased mean placental flow speed (Figure 5(f)); however, while the dependence on vein-to-artery ratio and villous density is relatively strong (metrics in Figures 5(a–d) span orders of magnitude), septal wall height variation has only limited influence on the organ-scale flow. We remark that the results in Figures 5(e,f) correspond to *N*_*V*_ */N*_*A*_ = 2 only (specifically, with two basal plate veins and one artery per cotyledon), hence the lack of variation.

The above trends are broadly reflected in the oxygen uptake, although with some interesting differences (Figure 6). Oxygen uptake decreases with *N*_*V*_ */N*_*A*_ (panel (a)), although with a marked ‘lag’, where mean uptake remains high (and approximately constant) across a significant range of *N*_*V*_ */N*_*A*_ values, before ‘short-circuiting’ of maternal flow between inlet and outlet flow pathways for large *N*_*V*_ */N*_*A*_ leads to a drop-off in uptake. While the dependence on villous density (through Da, panel (b)) follows the same increasing trend as shown in Figure 5(e), the dependence on septal wall height (panel (c)) is more complex, although we note that the influence of each of these aspects on uptake is weak – the variation in uptake shown in panel (b) is less than 30%, while it is less than 1% in panel (c). Figure 6(d) shows that oxygen uptake increases linearly with uptake rate (embodied by the Damköhler number, Dm), until the plateau occurring when this rate is sufficently high to extract all available solute from the incoming blood.

These extensive 2D results highlight the crucial importance of the vein-to-artery ratio, *N*_*V*_ */N*_*A*_, and villous density, characterised by Da and Dm, in influencing the IVS flow and oxygen uptake. Interestingly, they indicate that these macroscopic markers are insensitive to septal wall height. However, the flow within the placenta is inherently 3D, due to its irregular structural distribution, and while the 2D representation is advantageous due to its computational tractability, it only partially captures this complex, multiscale flow. Therefore, we use these 2D results to guide us in the more computationally intensive 3D problem, providing a comparison which also indicates the applicability or otherwise of such simplified descriptions. Further detail on the 2D problem setup along with the full presentation of simulation results can be found at [30].

### 3D Computational Study

Here, we build on the previous results to investigate in more detail the flow and transport dynamics in our physiologically inspired 3D placental domain, under variation both of its geometrical structure, and the villous density (Da) and uptake dynamics (Dm). The methodology for constructing our simulation geometries is described in Section **Placental Geometry**, but we note for clarity that we employ *ex vivo* data to define the macroscopic lobe structure, with lobule wall position and height, and the number and positions of arteries and veins being randomly allocated for each realisation.

The delivered placentas used are shown in **S1 Parameters and Geometry Description**, and indicated with red, blue and green markers in the figures in this section. Overall our 3D study comprises 54 unique geometries, upon which we have computed 298 flow and 3443 oxygen transport simulations. The coverage of the geometry and parameter spaces is not uniform, in part due to 2D results from **Preliminary 2D Simulation Study** and physiological considerations. Further detail on the simulation realisations and additional supporting plots can be found in **S4 Simulation Realisations and Supporting Plots**.

In the following figures, simulations using the rheological parameter values from **S1 Parameters and Geometry Description** (**S1 Table 1**) are referred to as having nominal parameter values.

### Placental Structure

#### Vein-to-artery ratio governs placental haemodynamics

In Figure 7 we investigate the influence of placental geometry, *N*_*V*_, *N*_*A*_, *N*_*V*_ */N*_*A*_ and Da on IVS flow in detail. We first examine the separate influence of arterial inflow and venous return on the flow dynamics in this 3D domain. Panel (a) shows little systematic dependence of average speed *S*_Ω_ on *N*_*V*_, in comparison to the much stronger correlation with *N*_*A*_ and *N*_*V*_ */N*_*A*_ (panels (b,c)), supporting the focus on *N*_*V*_ */N*_*A*_ in our 2D results presented

**Fig 7.**
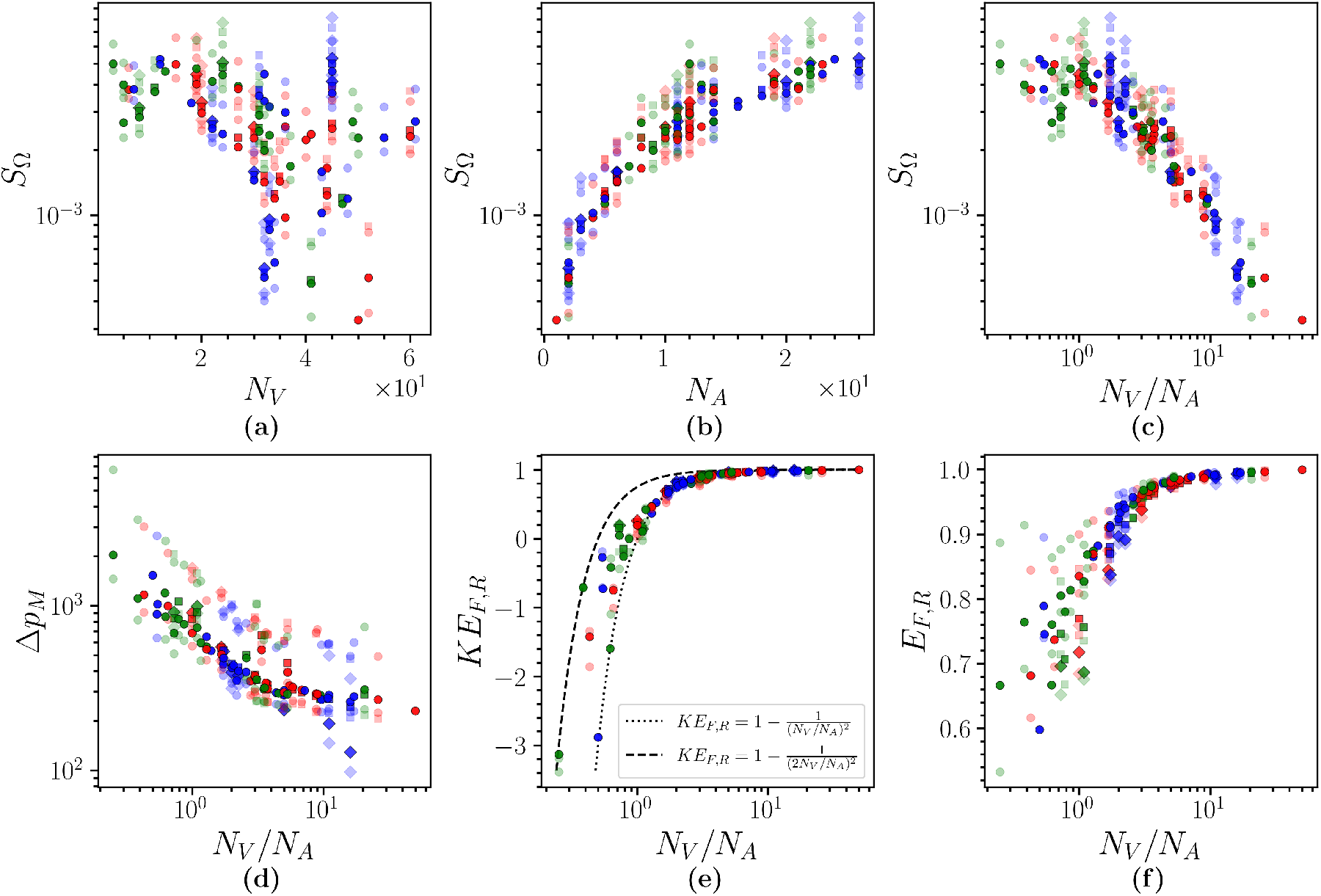
The variation of placental haemodynamics with vasculature. (a), (b), (c) Average fluid speed against number of veins, arteries and vein-to-artery ratio, respectively; (d) Maximum mean arterial pressure drop, (e) Kinetic energy flux loss ratio, (f) Total energy flux loss ratio, against vein-to-artery ratio. In (e), approximate laws for the behaviour of the kinetic energy flux markers against simulated values are shown by the dotted (*R*_*V*_ = *R*_*A*_) and dashed 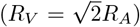 lines. Each scatter point corresponds to a different random geometry and the colour indicates the cotyledon configuration it uses. The circle, square and diamond markers represent data with Re = 4.54 *×* 10^2^, 9.08 *×* 10^2^ (nominal), 1.36 *×* 10^3^, respectively. Solid markers represent data with Da = 10^−4^ (nominal) while transparent markers represent data with Da = 10^−5^ or 10^−3^. above. Moreover, for the remaining flow markers (Δ*p*_*M*_, *KE*_*F,R*_, *E*_*F,R*_), coherent correlation is restricted to *N*_*V*_ */N*_*A*_, with similar levels of variation across realisations observed in both *N*_*V*_ and *N*_*A*_ (see **S4 Fig 2**); as such, discussion of the haemodynamic markers is henceforth largely focussed on the vein-to-artery ratio, *N*_*V*_ */N*_*A*_.

For a given vein-to-artery ratio and choice of Da, panels (c) and (d) of Figure 7 show that significant variation in average blood flow speed and pressure drop is associated with both macroscopic cotyledon structure and individual realisations, in which septal wall height and arterial/venous placement significantly influence local flow characteristics (see Figure 2 for an example of the complex flow patterns arising in the IVS). As in the above 2D results, we observe a clear decrease in average blood flow speed (panel (c)) with vein-to-artery fraction *N*_*V*_ */N*_*A*_ across the different geometries employed. Correspondingly, from panels (e) and (f) of Figure 7 we see increases in kinetic and total energy loss ratios.

Previous results indicate a significant influence of placental geometry on macroscopic IVS flow. Figure 7(e) compares these results against the approximate law for the dependence of the kinetic energy flux on the vein-to-artery ratio (32) obtained without appeal to the details of the placental geometry. This relationship is plotted in the special cases for which (i) *R*_*V*_ = *R*_*A*_ and (ii) 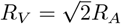. The significance of these choices is that in our model basal veins, septal veins and arteries have identical radii, while peripheral veins are larger (by a factor of 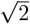). The close agreement between the approximate law and data, particularly for *N*_*V*_ */N*_*A*_ ≥ 1, indicates that this ratio strongly influences the kinetic energy flux loss ratio. Hence, dissipation of the kinetic energy in the placenta is primarily due to maternal vasculature rather than internal structure. Moreover, when *N*_*V*_ */N*_*A*_ is small the number of veins is typically small and the presence of peripheral veins (which have a significantly larger radius) has a much more prominent effect. This is clearly visible in Figure 7(e), wherein points corresponding to geometries that only (or predominantly) have outflow through peripheral veins appear on the dashed line, the approximate law for 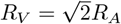. The larger vein radius corresponds to higher *KE*_*F,R*_: a similar effect is seen if the outflow area is increased by increasing the number of veins, and hence *N*_*V*_ */N*_*A*_. The associated box plot of the data from Figure 7 is shown in **S4 Fig 1**.

To further examine the slow flow environment, characteristic of IVS flow *in vivo*, we compare the mean placental flow speed obtained in each placental geometry realisation against the slow flow threshold (5 × 10^−4^ m s^−1^) identified in [5]. We found there is a statistically significant difference in *N*_*V*_ */N*_*A*_ for placental geometries which exhibit a slow flow environment, *i*.*e*. with mean placental flow speed below threshold, as determined by the Mann–Whitney *U* test (*U* = 1.27 *×* 10^4^, *p <* 1 *×* 10^−3^, median *N*_*V*_ */N*_*A*_ = 3.56 (slow set), 1.09 (fast set); *n* = 121). That is, the placental geometries exhibiting slow flow have on average a larger vein-to-artery ratio than those exhibiting higher speed, see Figure 8 for a box plot of the data. The statistically significant difference between the medians indicates that the vasculature strongly influences whether the IVS experiences preferential slow flow. Importantly, these results provide estimates for which values of *N*_*V*_ */N*_*A*_ may suggest potentially inadequate placental haemodynamics and hence indicate which pregnancies may be at risk of complications.

**Fig 8.**
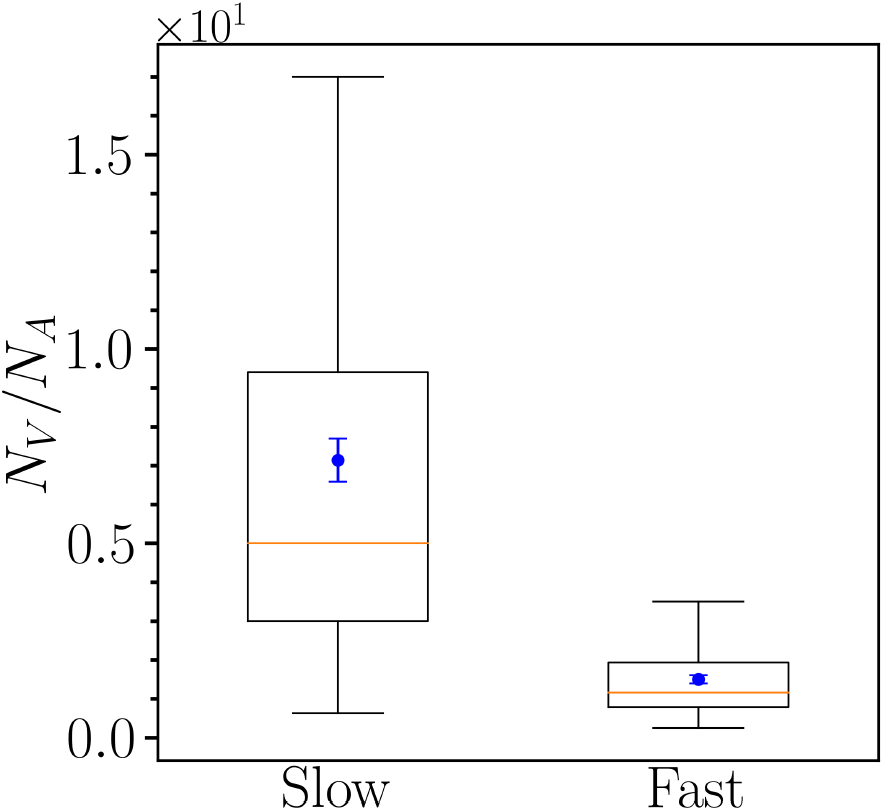
IVS slow flow. Box plot for flow solutions which have been segregated into two groups based on whether the mean placental speed, *S*_Ω_, is below or above the slow flow threshold value *V*_thresh_ = 5 *×* 10^−4^ m s^−1^, denoted by slow and fast, respectively. The independent variable of interest is the vein-to-artery ratio, *N*_*V*_ */N*_*A*_. The box plots show the quartile spread with the lower and upper box bounds representing the 25^th^ and 75^th^ percentiles and the yellow line the data median. Inside each box plot, the blue points represent the mean and the error bars are the standard error. Data used here corresponds to parameter values Re = 4.54 *×* 10^2^, 9.08 *×* 10^2^ (nominal) or 1.36 *×* 10^3^ and Da = 10^−5^, 10^−4^ (nominal) or 10^−3^.

#### Optimality of oxygen uptake relies on a cost-benefit balance between the slow flow regime and placental uptake efficiency

We now examine in detail the consequences of the previously studied IVS flow for oxygen delivery to the fetus in Figure 9. We first return to the dependence on arterial supply and venous return, considering how the mean uptake *O*_Ω,up_ varies with the number of each individual vessel type, as well as the total vein-to-artery ratio. Panels (a)–(d) reinforce previous results, indicating only a weak systematic dependence of uptake on the vein number, both from the point of view of each vessel type, and of overall number *N*_*V*_ . Considering panel (d) in particular, we see a slight decrease in uptake with number of veins overall, and a more significant reduction for *N*_*V*_ ≳ 30. We additionally see significantly larger variation associated with cotyledon geometry and individual realisations for larger values of *N*_*V*_ and/or higher Dm. Interestingly, panels (b) and (c) show little-to-no systematic dependence on either peripheral or septal wall vein number. These return pathways induce significant variation in uptake at intermediate numbers, while the variation in uptake associated with basal veins is increased for larger vein number.

**Fig 9.**
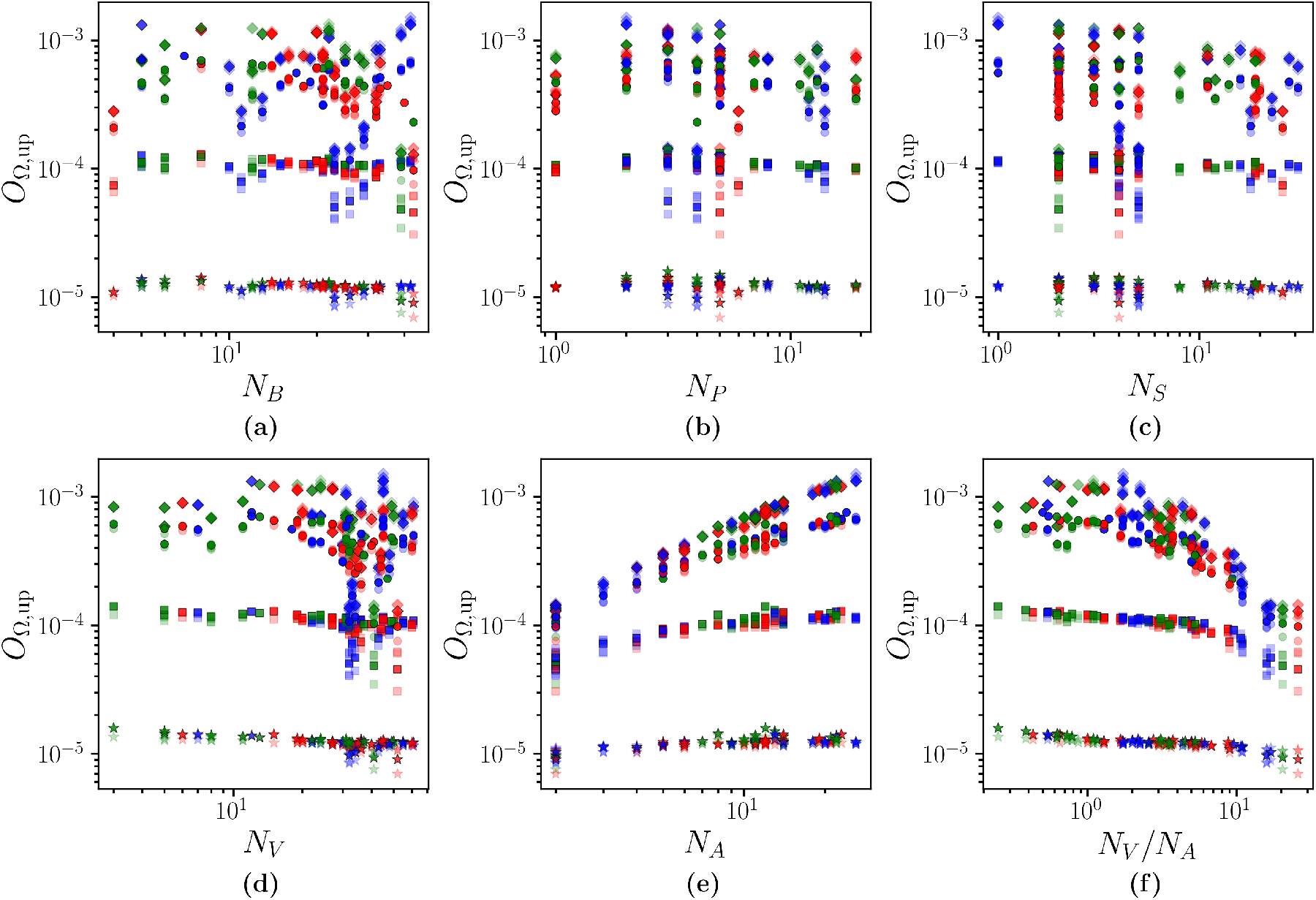
Oxygen uptake. Dependence of mean oxygen uptake *O*_Ω,up_ on the number of each vein type (basal plate *N*_*B*_, peripheral *N*_*P*_ and septal *N*_*S*_; total number *N*_*V*_), the number of arteries *N*_*A*_, and the total vein-to-artery ratio *N*_*V*_ */N*_*A*_. Each scatter point corresponds to a different random geometry and the colour indicates the cotyledon structure it has. Data with Dm = 1.11 *×* 10^−5^, 1.11 *×* 10^−4^, 1.11 *×* 10^−3^ (nominal), 1.11 *×* 10^−2^ are shown by the star, square, circle and diamond markers, respectively. Solid markers represent data with Re = 4.54 *×* 10^2^ (nominal), Da = 10^−4^ (nominal) while transparent markers represent data with Re = 4.54 *×* 10^2^ or 9.08 *×* 10^2^ and Da = 10^−5^ or 10^−3^.

The systematic dependence on artery number is significantly stronger, with an approximately linear increase in uptake with *N*_*A*_ (panel (e)), for sufficiently large Dm. Indeed the dependence on Dm shown in panel (e) reflects the analysis presented in Section **Reduction of the Exchange Model**: for increasing Dm we move from exchange-limited transport (independent of *N*_*A*_ at low Dm) to maternal-flow-limited transport at large Dm, in which *O*_Ω,up_ increases linearly with *N*_*A*_, as predicted by the reduced model presented in (18); for Dm ≫ Dm_*g*_, *O*_Ω,up_ ≈ Dm_*g*_, where Dm_*g*_ is defined in (17). The combined effect of artery and venous number on uptake, as expressed by the ratio *N*_*V*_ */N*_*A*_ (panel (f)), follows the trend observed in 2D, with efficient oxygen uptake across a wide range of *N*_*V*_ */N*_*A*_ values associated with the preferential slow flow environment (identified in Figure 7) before rapid drop-off for *N*_*V*_ */N*_*A*_ ≳ *O*(10). We remark that deviation from these trends is minimal across placental geometries with nominal structural parameter choices.

Figures 7 and 9(f) highlight a significant relationship in placental haemodynamics. A greater vein-to-artery ratio is vital for establishing a slow flow, low pressure environment, to avoid damage to the villi and hence the exchange pathway between maternal and fetal blood. However, it simultaneously results in the placenta becoming increasingly inefficient in delivering oxygen to the fetus. This cost-benefit behaviour suggests that an excess of venous return pathways (relative to arterial inflows) provides the ‘short-circuit’ effect that allows maternal blood to prematurely exit the placenta, thereby avoiding perfusion throughout the IVS and significantly hindering the ability for the villi to extract oxygen from the maternal blood.

We remark that increasing the number of basal plate veins has previously been observed to result in decreased oxygen uptake in lobule-specific simulations [31]. By providing the maternal blood with more venous return pathways close to the artery, the oxygenated blood can prematurely exit the lobule, insufficiently perfusing throughout the lobule and resulting in deoxygenated stagnation regions. The associated box plot of the data from Figure 9 is shown in **S4 Fig 4**.

#### Septal walls have a minimal impact on macroscopic haemodynamics

In Figure 10, we plot performance markers against mean cotyledon wall height, *H*_*c*_. Similar to the results obtained in 2D (results presented in Figures 5 and 6) we observe no strong correlation between the macroscopic performance markers and *H*_*c*_. This somewhat surprising result is in line with that presented in Figure 9 which shows no systematic dependence of uptake on the number of septal veins. We do note, however, that changes in septal wall height induces very significant variation in flow and uptake markers (*cf*. Fig. 9). Hence, while local flow dynamics is certainly influenced by these small scale structural features, their detail does not appear to influence macroscopic placental function in a predictable manner.

**Fig 10.**
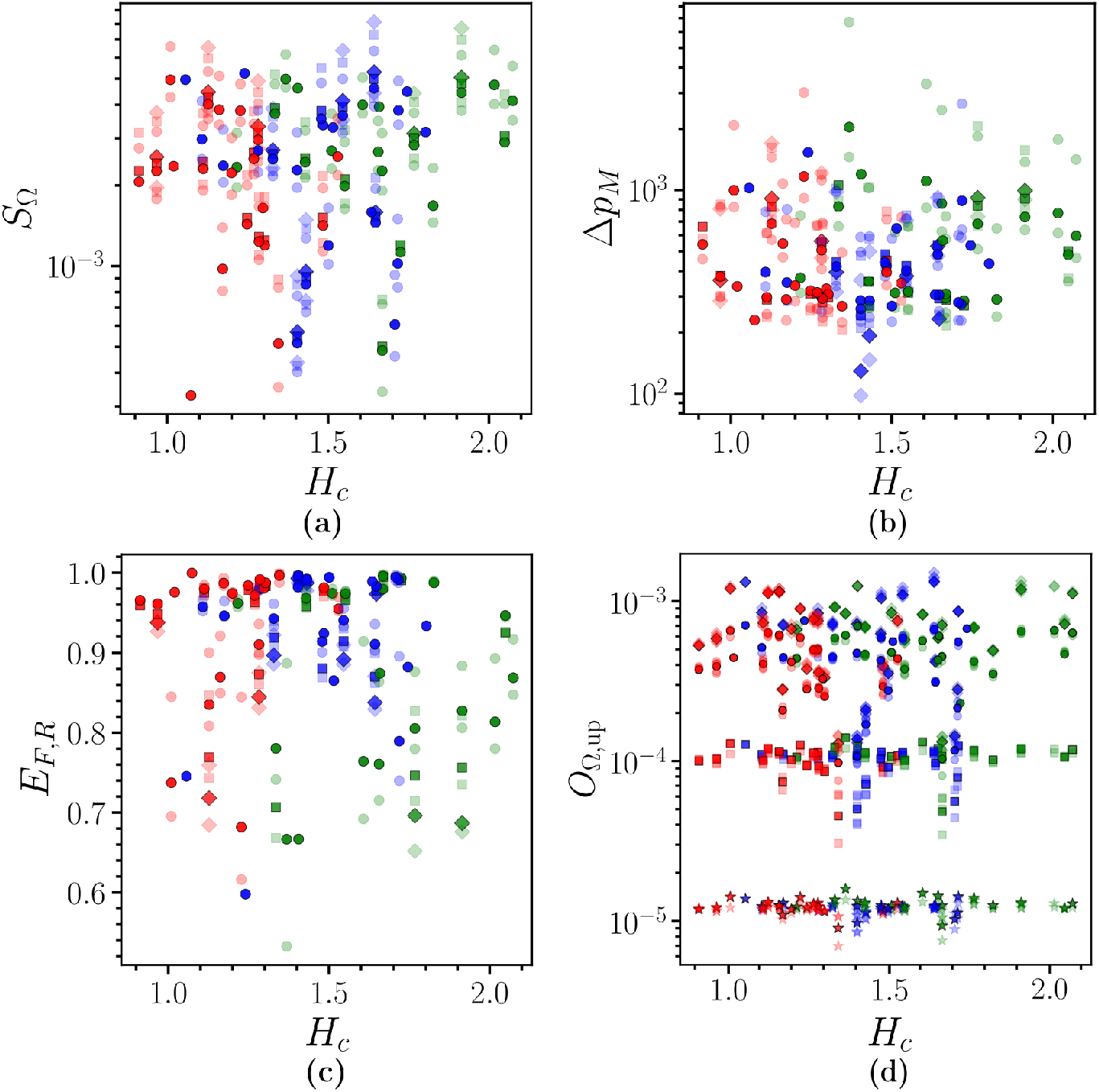
Dependence of IVS flow and oxygen uptake on septal walls. (a) Average fluid speed, (b) maximum mean arterial pressure drop, (c) total energy flux loss ratio, (d) mean oxygen uptake presented as a function of average cotyledon wall height. Each scatter point corresponds to a different random geometry and the colour indicates the cotyledon structure it has. For panels (a), (b) and (c), the circle, square and diamond markers represent data with Re = 4.54 *×* 10^2^ (nominal), 9.08 *×* 10^2^, 1.36 *×* 10^3^, respectively. Solid markers represent data with Da = 10^−4^ (nominal) while transparent markers are data with Da = 10^−3^ or 10^−5^. For panel (d), star, square, circle and diamond markers correspond to Dm = 1.11 *×* 10^−5^, 1.11 *×* 10^−4^, 1.11 *×* 10^−3^, 1.11 *×* 10^−2^, respectively.

#### Villous density regulates the IVS flow environment

In Figure 11, we present plots of various performance markers versus changes in the IVS villous density, characterised by Da. We see in panels (a) and (b) that an increase in Da corresponds to increased average speed and reduced pressure drop, respectively. This is expected as an increase in Da corresponds to a higher permeability of the IVS, which is associated with reduced resistance to maternal flow and therefore smaller inertial losses between the CC and IVS. In panels (c) and (d), while we see that that Da has most significant influence on *KE*_*F,R*_ and *E*_*F,R*_ at small *N*_*V*_ */N*_*A*_, the behaviour is influenced by *N*_*V*_ */N*_*A*_ for the full range of Da tested. Interestingly, for a given value of *N*_*V*_ */N*_*A*_, *KE*_*F,R*_ increases with increasing Da, suggesting that higher inertia in the IVS is responsible for greater kinetic energy flux losses than higher resistance of the IVS. We note that the negative values of *KE*_*F,R*_ in Fig. 11(c) arise in the physically unlikely regimes for which *N*_*V*_ ≲ *N*_*A*_ (see also Fig. 7(e) and equation (32)).

**Fig 11.**
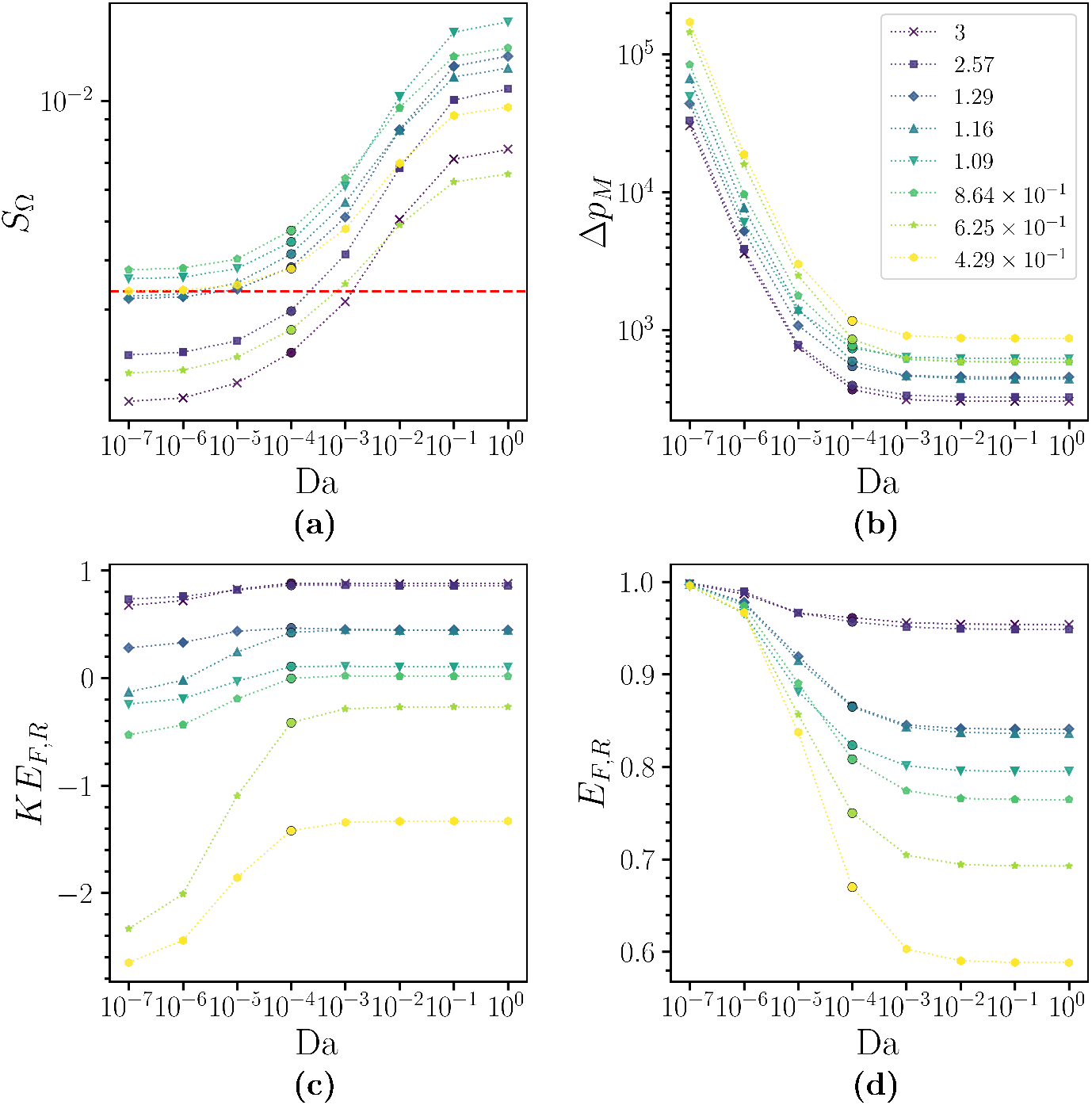
Villous density and IVS haemodynamics. (a) Average fluid speed, *S*_Ω_, (b) maximum mean arterial pressure drop, Δ*p*_*M*_, (c) kinetic energy flux loss ratio, *KE*_*F,R*_, (d) energy flux loss ratio, *E*_*F,R*_, plotted against Da. The vein-to-artery ratio, *N*_*V*_ */N*_*A*_, for each plotted geometry is shown in the legend in panel (b). The circle markers represent data from flow problems with nominal parameter values, the other markers distinguish the different *N*_*V*_ */N*_*A*_ data points. In panel (a), the red dashed line represents the slow flow threshold value.

Regarding trends with Da, we notice all markers begin to plateau for large Da, with this behaviour delayed for *S*_Ω_ relative to the other markers. This plateau is expected as Da →𝒪 (Re^−1^), since inertial effects begin to dominate the flow dynamics. For Da ≫𝒪 (Re^−1^), the permeability of the IVS only acts as a perturbative effect on flow dynamics. A noticeable trend is seen in panels (b), (c) and (d) where, for a given Da, monotonic behaviour of the performance markers occurs with varying *N*_*V*_ */N*_*A*_ for small Da. However, this is not seen in panel (a), and we also notice in panels (a) and (b) that the ordering of the geometries changes as Da increases. These can be explained via two different mechanisms for small and large Da.

For small Da, the flow within the IVS is dominated by the viscous drag encoded in the Darcy term and the maternal blood preferentially follows paths of least viscous dissipation. Following a similar venous return pathways argument used to explain Figure 9, it is likely that in this regime the vessels’ locations can influence how well, and how fast, the maternal blood perfuses throughout the placenta. While *N*_*V*_ */N*_*A*_ has a large influence on *S*_Ω_ (Figure 7), the non-monotonic behaviour of the geometries’ ordering with *N*_*V*_ */N*_*A*_ suggests that there are additional structural factors which influence this. For small Da, large resistances of the IVS results in monotonic behaviour of the pressure drop and energy markers with *N*_*V*_ */N*_*A*_. For large Da, the flow becomes increasingly dominated by inertial behaviour and governed by the non-linear advection term. It is, however, still being influenced strongly by the vessels’ locations, and the variability between geometries in local inertial flow patterns weakens the general patterns visible in the observed behaviours for small Da.

While Figure 11 highlights the potential for favourable haemodynamic regimes when the IVS behaves as a dense porous medium, we only obtain partial insight into the flow within the IVS. Specifically, the question of where the flow is expending its energy within the placenta arises: the central cavity or the IVS. In view of the discussion in **Introduction** on energy expenditure in the IVS, we further investigate these performance markers against viscous dissipation rate in the IVS, 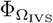, in Figure 12. In panels (a) and (b), we see there are generally smaller mean placental speeds and pressure gradients for smaller 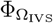. Importantly, panels (c) and (d) indicate that a lower 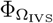 suggests greater kinetic energy and total energy loss ratios across the placenta, reinforcing the role that the IVS plays in healthy placentas to transform the inertial spiral artery flow into a viscous creeping flow. Moreover, via 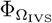, this viscous creeping flow coincides with a reduction of stress exerted on the villi in the IVS.

**Fig 12.**
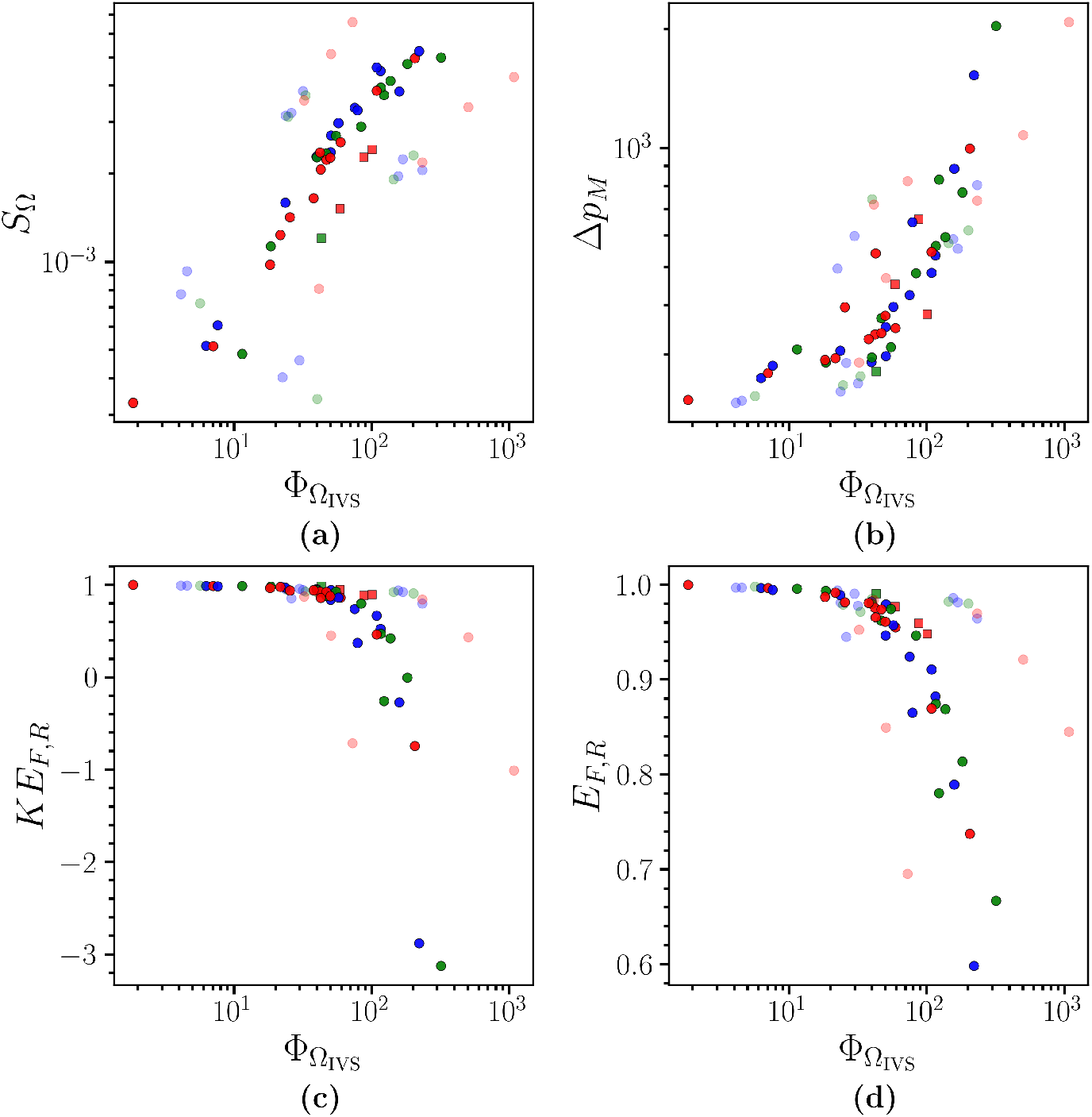
IVS haemodynamics and viscous dissipation rate. (a) Average fluid speed, *S*_Ω_, (b) maximum mean arterial pressure drop, Δ*p*_*M*_, (c) kinetic energy flux loss ratio, *KE*_*F,R*_, (d) energy flux loss ratio, *E*_*F,R*_, plotted against 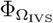. Each scatter point corresponds to a different random geometry and the colour indicates the cotyledon configuration it uses. The circle and square markers represent data with Re = 4.54 *×* 10^2^ (nominal), 9.08 *×* 10^2^, respectively. Solid markers represent data with Da = 10^−4^ (nominal) while transparent markers are data with Da = 10^−3^ or 10^−5^.

#### Maternal-fetal exchange highlights redundancies in placental function and short-circuiting effects

In Figure 13 we compare the volume-averaged oxygenation of each cotyledon and the whole placenta. In panel (a), we observe the expected monotone increase in oxygen uptake with Dm in each separate cotyledon (crosses) and overall (yellow circles). Different behaviour is seen in panel (b), where one cotyledon exhibits a rapid decrease in uptake once Dm is large enough. This is due to the lack of self-sufficient blood supply in this particular geometry cotyledon: the cotyledon does not contain an artery. Therefore, while fresh blood is still transferred into it from neighbouring cotyledons to its venous return outlets, once the uptake rate exceeds a given threshold, all the available oxygen in this blood has already been transferred to the fetal vasculature, creating a hypoxic environment within this cotyledon.

**Fig 13.**
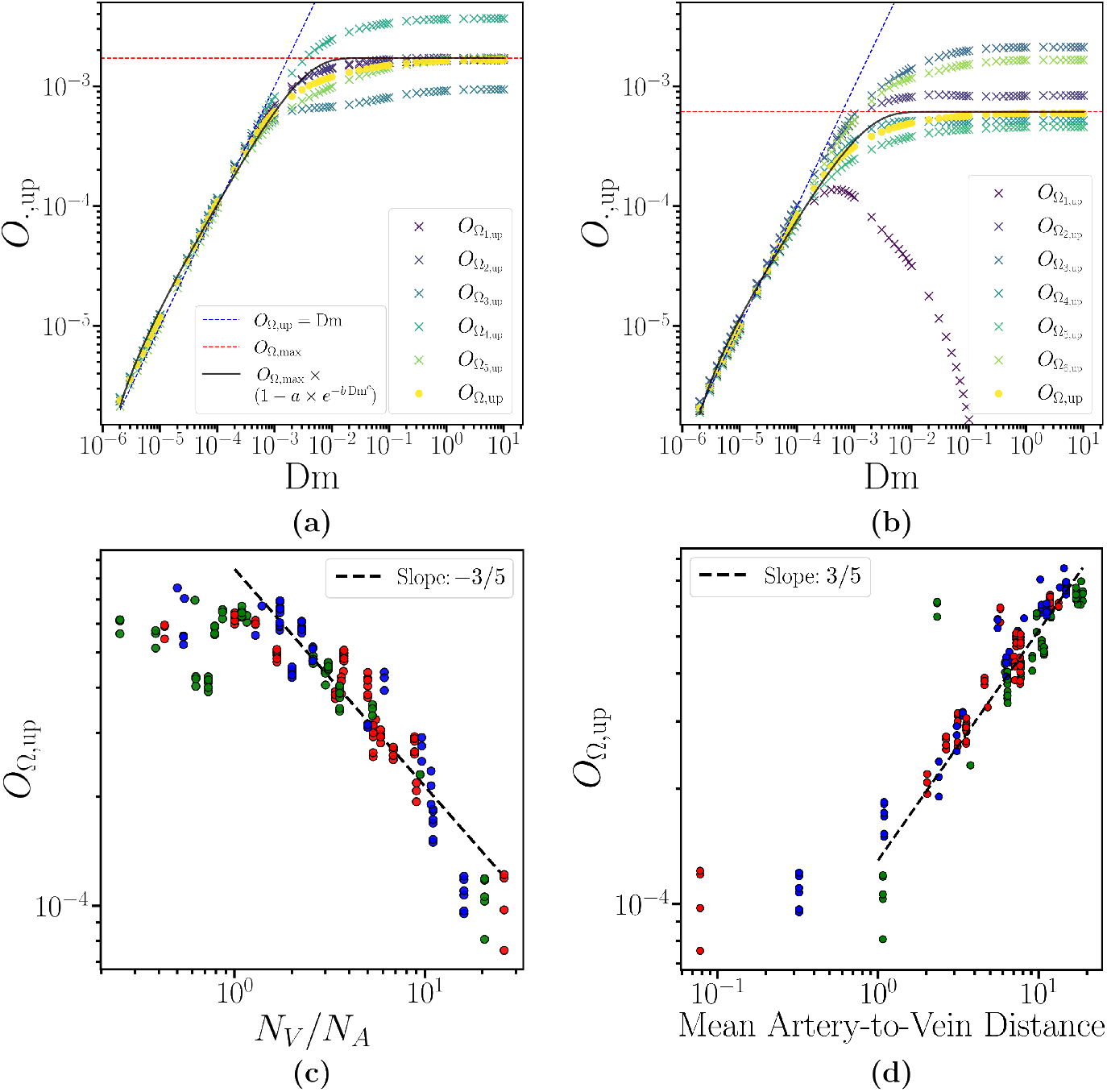
(a) II, *N*_*V*_ */N*_*A*_ = 8.63 *×* 10^−1^, (b) I, *N*_*V*_ */N*_*A*_ = 5.5 show mean oxygen uptake, 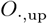, against Dm for two different geometries, where the roman numerals indicate the cotyledon configuration used from **S1 Parameters and Geometry Description**. Crosses indicate uptake in each cotyledon making up the geometry, and the yellow circles the total uptake. The blue and red dashed lines correspond to the fetal uptake-limited (linear) and maternal flow-limited (35) regimes, respectively. The black line represents the best fit of the exponential function (36) to *O*_Ω,up_, with the exact form shown in the legend of (a); fitted values are given in **S4 Simulation Realisations and Supporting Plots**. Panels (c), (d) show *O*_Ω,up_ against vein-to-artery ratio, *N*_*V*_ */N*_*A*_, and mean artery-to-vein-distance, respectively. The colours indicate the different cotyledon configurations I–III, as previously. Scatter points in: panels (a), (b) correspond to parameter values Re = 4.54 *×* 10^2^ (nominal) and Da = 10^−4^ (nominal); in panels (c), (d) correspond to parameter values Re = 4.54 *×* 10^2^ (nominal) or 9.08 *×* 10^2^, Da = 10^−5^, 10^−4^ (nominal) or 10^−3^ and Dm = 1.11 *×* 10^−3^ (nominal). The dashed lines represent the relationship *O*_Ω,up_ ∝ (*N*_*V*_ */N*_*A*_)^−3*/*5^ ∼ (mean artery-to-vein distance)^−1^ obtained from **S6 Uptake in a half-space** with the approximation 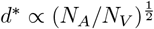.

In the small-Dm regime, we see that the limiting factor in the uptake is the rate of uptake and there is a linear relationship between Dm and 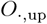 (highlighted by the blue dashed line in each figure). In contrast, in the high-Dm regime the uptake is limited by the oxygen supply; the upper limit is hence

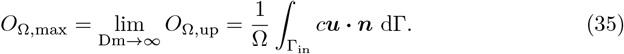

This limit is shown by the horizontal dashed red line in panels (a) and (b). We note further that despite the deficient cotyledon discussed above, the *overall* exchange function of the placenta in panel (b) is largely unaffected, highlighting a form of ‘redundancy’ in design that supports organ function even in the presence of failure of its individual components.

We now return explicitly to the fetal coupling examined in **Reduction of the Exchange Model**. Figures 13(a,b) identify Dm_*g*_ from equation (17) as the intersection between the dashed red and blue lines. Taking into account the limiting behaviour and exponential-like dependence evident in Figs 13(a,b), and the asymptotic behaviour identified in (18), we propose the ansatz

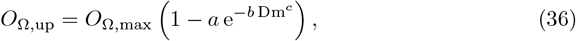

with fitting parameters *a, b* and *c*, and we note that *a* = 1, *b* = 1*/*Dm_*g*_ and *c* = 1 correspond to the asymptotes from (18). For the geometrically complex simulations carried out here, the additional generality provided by (36) allows for a better fit to the simulated data in the range of maternal Damköhler numbers relevant to real placenta flows.

In panels (a) and (b), the resulting fit is shown by the black line showing that the dependence of uptake on Dm is well-captured in general. We note, however, that the intermediate region between the low- and high-Dm regimes is not fully captured, and that the quality of fit varies between panels (a) and (b). Furthermore, the fitted values of *a, b* and *c* do not conform precisely to the theoretical values noted above (values provided in **S4 Fig 5**). Together these observations suggest that the complexity of the flow and transport imbued by the details of the placental geometry result in some small deviation from the theoretical estimates. While detailed investigation is out of the scope of this work, in **S4 Fig 5** we additionally plot the fitting function parameters against *N*_*V*_ */N*_*A*_ for a range of geometries, observing distinct correlations with all three. This indicates that consideration of the vasculature may play a key role in improving the form of (36).

As exposed in [24] and discussed above, a potential mechanism for placental inefficiency is through ‘short-circuiting’ behaviour, whereby proximity between arterial supply and venous drainage can enable solute to leave the IVS before being uptaken by the fetal vasculature. To illustrate the way in which such short-circuiting reduces solute exchange, in **S6 Uptake in a half-space** we consider an idealised colinear arrangement of an artery between two identical veins. The artery supplies a volume flux 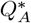, carrying solute with concentration 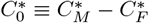 and the artery-vein distance is *d*^*^. Provided the solute uptake rate 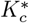 is weak 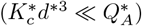, exchange takes place far enough from the artery and vein for the flow to be approximately dipolar.

Exploiting the self-similar form of dipolar streamlines leads to the approximation for the exchange rate as 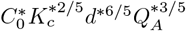(a numerical prefactor is given in (83)).

In Figure 13(c,d) we present the dependence of *O*_Ω,up_ on *N*_*V*_ */N*_*A*_ (partial reproduction of Figure 9(f)) and on the mean artery-to-vein distance, respectively. We emphasise that in these simulations, Dm is fixed at its nominal value, while the simulation geometries (indicated by colour) and other parameters (not indicated) are varied: the data points here are the collection of simulations for Re = 4.54 *×* 10^2^, 9.08 *×* 10^2^, Da = 10^−5^, 10^−4^, 10^−3^ with Dm = 1.11 *×* 10^−3^.

The complexity of our 3D simulation geometry means that there is not a direct equivalence to the idealised model from Section **Reduction of the Exchange Model**; however, associating *d*^*^ ∝ (*N*_*A*_*/N*_*V*_)^1*/*2^ (*i*.*e*. estimating artery-to-vein distance from the area density of veins and arteries) we expect *O*_Ω,up_ ∝ (*N*_*V*_ */N*_*A*_)^−3*/*5^. Likewise, we expect the inverse relationship with mean distance. The dashed lines in panels (c,d) confirm this. Indeed, it is clear that, as indicated by Section **Reduction of the Exchange Model**, this correlation is largely insensitive to significant variation in parameters and geometrical detail. It is noteworthy that the correspondence of our 3D simulation results does not only apply for extreme choices—the agreement remains excellent for values of *N*_*V*_ */N*_*A*_ ≈ 2 − 3, *i*.*e*., in the physiological range identified earlier. Our simulations thereby indicate that this short-circuiting effect may be of relevance in actual placentas.

Finally, we remark that the corresponding short-circuiting effect in the flow field is also evident in the dropoff in *S*_Ω_ shown in Figure 7(c): by the same arguments as above, the volume of perfused IVS will scale with the cube of source-sink distance, which decreases with *N*_*V*_ */N*_*A*_. Equivalent observations can be made in Figure 5(a).

## Conclusions and discussion

In this study, we identify links between placental structure, maternal flow haemodynamics and solute transport. By combining comprehensive 2D and 3D simulation of an *ex vivo* data-informed mathematical model with analytical model reduction approaches, we have highlighted the key features of placental structure that can influence and regulate the flow within the placenta.

A key finding of this work is that the vein-to-artery ratio, *N*_*V*_ */N*_*A*_, is an important determinant of the flow haemodynamics and fetal oxygen uptake. We show that this structural parameter is largely responsible for the cost-versus-benefit relationship of maintaining a slow, low pressure flow environment in the IVS while maximising fetal oxygen uptake. This complements current understanding by tying maternal inflow to venous return in the placenta, showing how the vasculature within the placenta can influence its function.

Current literature associates optimal placental function with a ‘slow-flow’ regime, in which the damaging viscous energy expenditure in the dense IVS is balanced against effective perfusion that supports efficient maternal-fetal solute exchange. Our investigations highlight how arterial supply and venous return are vital components in regulating this balance. Notably, we determine that for *N*_*V*_ */N*_*A*_ ≈ 3, the mean placental speed switches from above to below the slow flow threshold specified in [5], and at which point we also see a reduction in the pressure gradients and relatively high oxygen uptake to the fetal vasculature. Our simulations have hence identified a physiologically relevant regime emphasising the cost-versus-benefit principle behind the structure of the placenta, wherein its structure is the result of an optimisation over its competing demands; *i*.*e*., there should be enough veins per artery to avoid a damaging, high speed, haemodynamic environment, but not so many as to inhibit effective solute transfer.

To understand this maternal-fetal transfer, we derive two reduced descriptions. The first applies a homogenisation argument to the solute transport problem, highlighting the transitions between exchange-limited transport, maternal-flow-limited transport and fetal-flow-limited transport as key model parameters are varied. In the present model, fetal conditions are captured parametrically via the dependence of maternal Damköhler number Dm on fetal Damköhler number Df, via (14). The second exploits an idealised source-sink description, in a suitable asymptotic regime, to indicate how uptake inefficiency arising from increased venous return pathways is driven by short-circuiting behaviour whereby maternal blood prematurely exits the placenta. In each case, good agreement between the analytical approximations and the simulation results is observed. Moreover, in the latter case, we demonstrate that the analysis of the short-circuiting effect remains accurate even for vein-and-artery numbers and placements that accord with physiological relevance, suggesting that this effect may be of significance in real placentas. To our knowledge no specific evidence of this, either in an *in vivo* or *ex vivo* setting, has been reported, motivating future experimental investigation.

The preferential slow flow environment of healthy placentas has been associated with lower stresses and viscous dissipation in the IVS [10]. Therefore, (relatively) low energy losses in the flow may suggest inadequate functioning of the placenta. The simulation results indicate a correlation between lower viscous dissipation in the IVS and higher placental energy losses. Additionally, sparser villi structures, manifesting in the haemodynamics via a decrease in permeability of the flow, may indicate poor placental health. Our simulations reinforce this point, highlighting the importance of IVS permeability in ensuring the maternal flow achieves a slow flow environment.

Current estimates of the numbers of arteries and veins (and indeed their type) vary widely, and are not characterised on an individual or group basis *in vivo*. Overall our study hence highlights that accurate characterisation of these flow pathways *in vivo* (such as via MRI or other non-invasive imaging modalities) may provide an effective approach to identify deficiencies in placental haemodynamics that are associated with adverse pregnancy outcome. This has implications in clinical practice: while current antenatal care focuses on determining maternal arterial inflow, our results demonstrate the ratio of arterial inflow to venous return regulates the haemodynamic function within the placenta. Therefore, we suggest that the venous return, or more specifically the vein-to-artery ratio, is a critical yet understudied biomarker for placental health. Future developments in non-invasive imaging techniques which characterise this ratio directly or indirectly may thereby aid clinical pregnancy care and monitoring.

A surprising observation from our simulation results is that the presence of septal walls (and their associated influence on routes for venous return) has no clear systematic impact on our measurements of macroscopic placental function, though we note that the impact that such variation will undoubtedly have on local flow paths is manifest in significantly increased variability in these markers across computational geometries. We emphasise that our focus has been on the impact of these various geometrical and model parameters on function at the global scale and so any more systematic effect may be blurred by cross-cotyledon transport, and a form of the redundancy that we have highlighted. Moreover, our choice to include basal plate veins (in accord with the extant literature, and especially those modelling studies noted earlier) would naturally help to lessen their specific effect. We defer to future work the influence of septa on more local flow and transport features, which such return routes will naturally affect. We also recognise that our choice of flow and uptake model is intentionally simplified. Important future work includes more detailed consideration of the flow behaviour in variable-porosity media, and more faithful models of solute uptake and transport that reflect the specifics of (for example) oxygen-haemoglobin binding.

## S1 Parameters and Geometry Description

### Parameters

In **S1 Table 1**, we present the rheological parameter values which define the flow problems and in **S1 Table 2**, the geometrical parameters which define the placental geometries. These values are explored to study their effect on the resulting haemodynamics.

The characteristic length 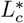 is chosen as a mesoscopic-like length scale which characterises the transition length between the small length scale associated with the arteries 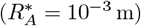 and the organ length scale, characterised by the chorionic plate radius 9.07 *×* 10^−2^ m. The characteristic velocity scale 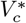 is obtained from the spatial average of the imposed Poiseuille arterial inlet velocity. In dimensional form the boundary condition is

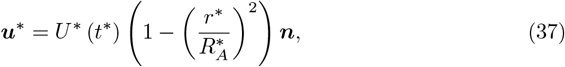

where *U* ^*^ (*t*^*^) denotes the pulsatile peak speed, obtained from Doppler ultrasound, scaled appropriately for the third trimester [32–34] and ***n*** is a unit inward-pointing normal to the boundary. We find 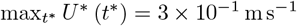, from which 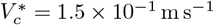.

The governing equations are expressed in dimensionless form by scaling variables on 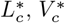 plus the characteristic time 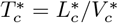 and pressure 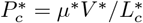. The resulting equations involve the dimensionless parameters Re, Da, Pe, Dm, as defined in Table 1.

We remark that in the simulations, a change in Re is implemented via a variation in 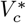, i.e. the peak arterial inflow speed, as we take 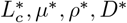 to be fixed, as indicated by a parameter range of N/A. Therefore, a variation in Pe also occurs in the corresponding transport simulation. In the text, to avoid unnecessary repetition, we only report on Re values as the corresponding Pe can be inferred using the nominal values from **S1 Table 1** and specified Re value(s).

### 3D Geometry

We approximate the shape of the placenta by a spherical cap with base radius *R* and height *h*. The apex of the spherical cap is taken as the origin, *O*, at ***r*** = **0**. The spherical and flat surface represents the basal and chorionic plate, respectively. The cotyledons are generated via a Voronoi tessellation which covers the spherical surface [29]. The lobules are generated as a centroidal Voronoi tessellation over each cotyledon by using Lloyd’s algorithm with a reasonably large error to ensure a degree of uniformity in the lobule sizes [45].

Cell centres must be provided to generate the Voronoi tessellations. To this end, we have taken images of delivered placentas provided by Nottingham University Hospitals and then manually marked the cotyledon septal walls. Due to the irregular shape of the *ex vivo* chorionic plate, we only considered a circular section of the image. We then used Matlab’s Image Processing Toolbox [46] to identify the cell centres of the cotyledons. In **S1 Fig 1**, we show three placentas with their cotyledon septal walls marked in blue. Overlapped are the idealistic geometries which used the cell centres of the cotyledons to generate the Voronoi tessellation which models the shape of the cotyledons.

For each lobule’s Voronoi tessellation, we assume that there are 4 or 5 lobules per cotyledon. Lloyd’s algorithm is used to generate the cell centres for each set of lobules from a set of initially randomly placed cell centres.

**S1 Table 1.**
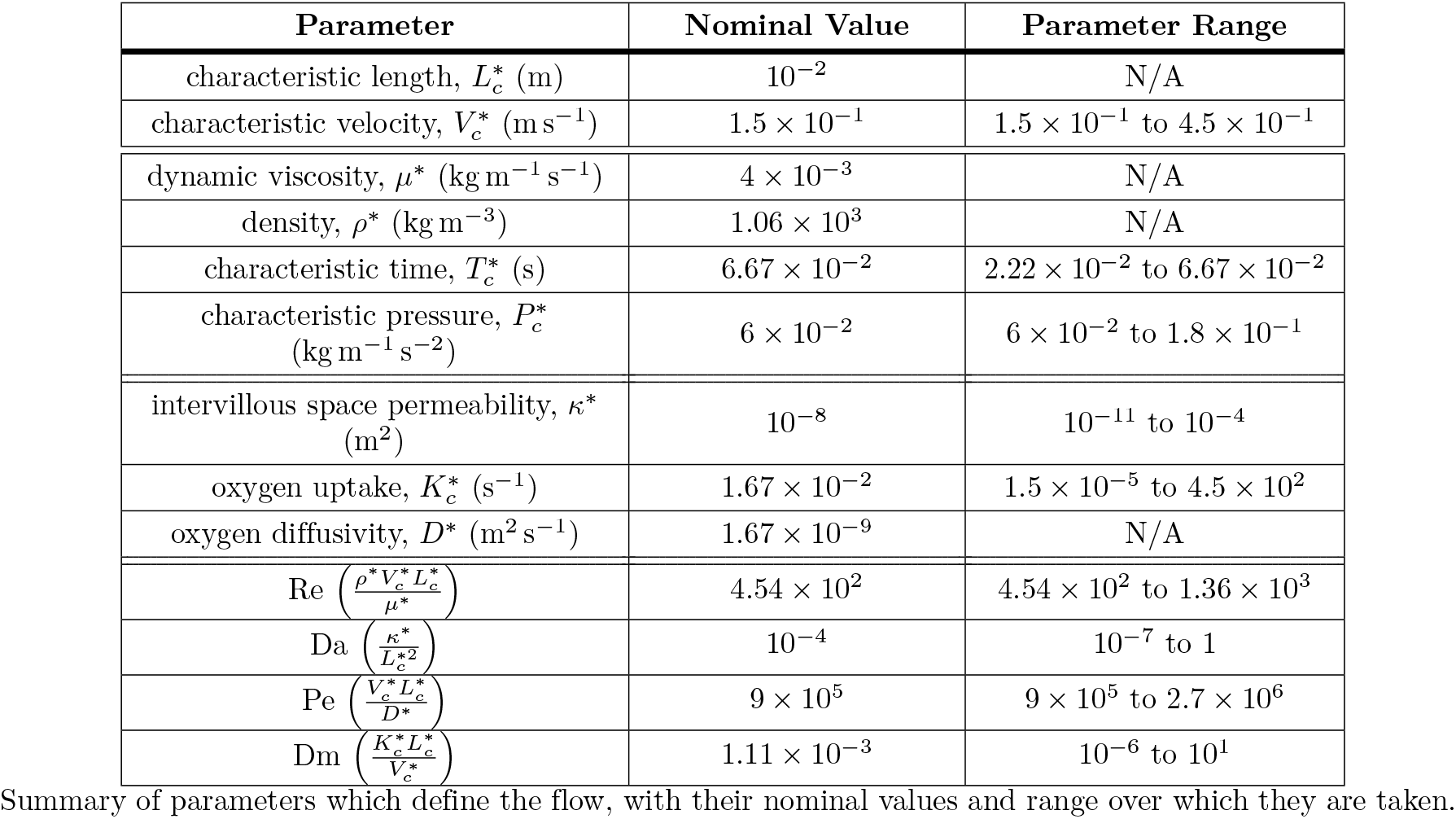
Rheological parameters.

**S1 Fig 1.**
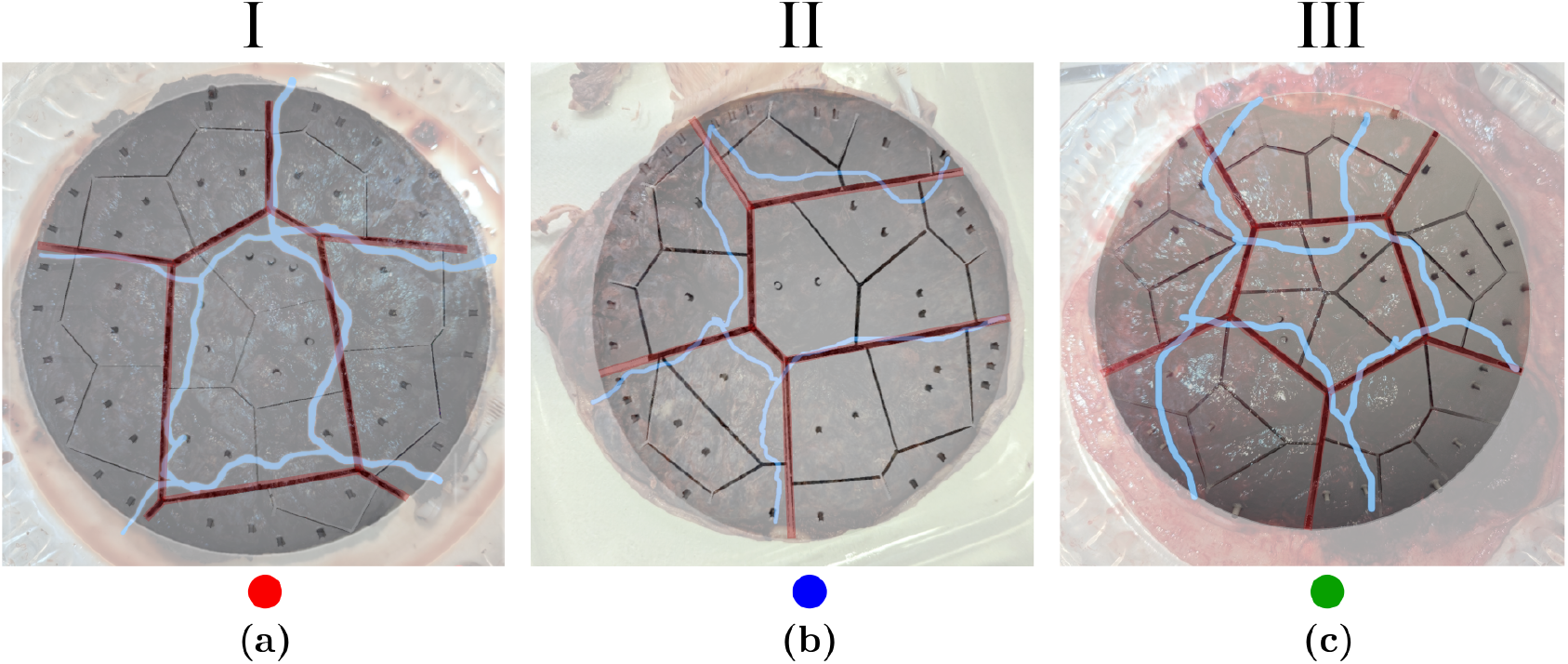
Data-informed cotyledon configurations. Three delivered placentas with the recreated Voronoi tessellated geometries imposed on top of each. The blue and red lines indicate the cotyledon septal walls of the placenta and the Voronoi cell borders, respectively. The numeral above each image denotes the cotyledon configuration, with the colour below corresponding to the scatter point colour of geometries using this configuration in figures of the main text.

**S1 Table 2.** Geometrical parameters.

| Parameter | Representative Value | Parameter Range |
| --- | --- | --- |
| chorionic plate radius (m) | $9.07 \times 10^{-2}$ | N/A |
| central placental height (m) | $3.15 \times 10^{-2}$ | N/A |
| placental volume (m<sup>3</sup>) | $4.28 \times 10^{-4}$ | N/A |
| central cotyledon wall height (m) | $1.4 \times 10^{-2}$ | $7 \times 10^{-3}$ to $2.1 \times 10^{-2}$ |
| central lobule wall height (m) | $7 \times 10^{-3}$ | $3.5 \times 10^{-3}$ to $1.5 \times 10^{-2}$ |
| cotyledon wall thickness (m) | $2 \times 10^{-3}$ | N/A |
| lobule wall thickness (m) | $1.3 \times 10^{-3}$ | N/A |
| cavity minor axis (m) | $4 \times 10^{-3}$ | N/A |
| cavity major axis (m) | $\frac{1}{3}$ of placental height<br>at artery location | N/A |
| spiral artery radius, $R_A^*$ (m) | $10^{-3}$ | N/A |
| basal plate vein radius, $R_B^*$ (m) | $10^{-3}$ | N/A |
| peripheral vein radius, $R_P^*$ (m) | $1.41 \times 10^{-3}$ | N/A |
| septal wall vein radius, $R_S^*$ (m) | $10^{-3}$ | N/A |
| no. cotyledons, $N_C$ | 6 | 5 to 6 |
| no. lobules, $N_L$ | 30 | $4N_C$ to $5N_C$ |
| no. spiral arteries, $N_A$ | 15 | 1 to $N_L$ |
| no. basal plate veins, $N_B$ | $N_L$ | 0 to $2N_L$ |
| no. peripheral sinus veins, $N_P$ | 6 | 0 to 15 |
| no. septal wall veins, $N_S$ | $N_C$ | 0 to $5N_C$ |
| typical no. mesh elements | $\sim 5 \times 10^5$ | $\sim 3 \times 10^5$ to $\sim 8 \times 10^5$ |

Summary of parameters which define the placental geometries with their representative values and range over which they are taken.

**S2 Fig 1.**
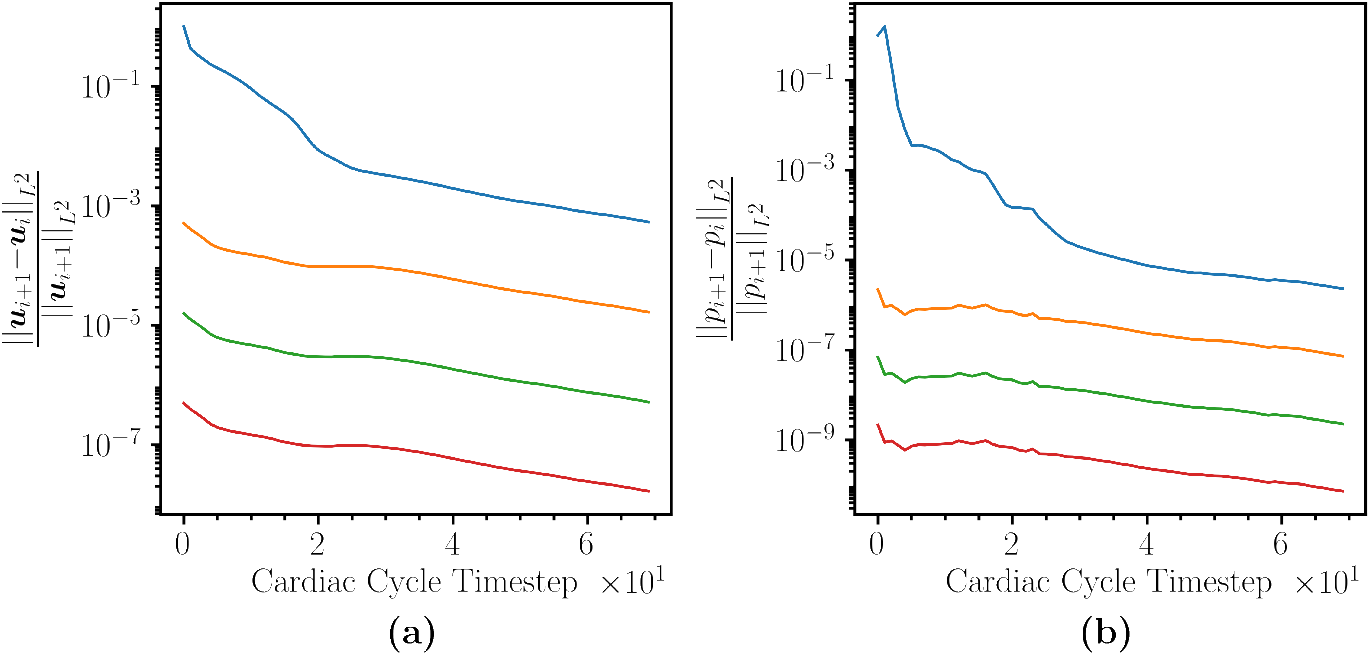
*L*^2^ norm relative convergence plots for (a) ***u***, (b) *p* across 5 cardiac cycles. Variable subscript *i* refers to the solution of cardiac cycle *i*.

In each cotyledon, we randomly place between 0–5 septal wall veins. In each lobule, we randomly assign 0 or 1 spiral arteries to the centre of the lobule and between 0–2 basal plate veins. The basal plate veins are placed 7.75 mm along the tangent to the spherical cap from the spiral artery’s centre. Along the periphery, we randomly place between 0–15 peripheral veins. We denote the total number of: spiral arteries by *N*_*A*_; veins by *N*_*V*_ ; basal plate veins by *N*_*B*_; peripheral veins by *N*_*P*_ ; septal wall veins by *N*_*S*_. In Figure 2, the interior of an idealised geometry with some of the vessel types highlighted can be seen in the second panel.

We denote the placenta’s domain by Ω, with the *N*_*C*_ cotyledon sub-domains denoted by Ω_*i*_, *i* = 1, …, *N*_*C*_ with 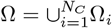. The cotyledon sub-domains are identified via their Voronoi tessellation cell.

## S2 Cardiac Cycle Convergence

In **S2 Fig 1**, we plot the normalised relative convergence under the *L*^2^ norm of the velocity and pressure solution for one simulation over successive cardiac cycles. In **S2 Table 1**, we present the normalised relative difference of the associated flow performance markers at successive cardiac cycles. We observe that after the first cardiac cycle, there is a distinct drop in the difference. This is primarily due to the transients present in the initial cardiac cycle becoming damped out by the subsequent cardiac cycles from the steady-like, viscous flow which dominates the majority of the flow within the placenta.

## S3 Homogenisation

### *µ*CT data acquisition

To obtain an estimate for IVS permeability, we employ a classical homogenisation approach, exploiting (synchrotron) *µ*CT data to define the relevant microscale geometry.

**S2 Table 1.** Cardiac cycle convergence of performance markers. Normalised relative difference of some flow performance markers, denoted by *J*, for one simulation at successive cardiac cycles, denoted by *i*.

| Quantity, $J$ | $\frac{J_{i+1}-J_i}{J_{i+1}}$ | | | |
| --- | --- | --- | --- | --- |
| | $i = 1$ | $i = 2$ | $i = 3$ | $i = 4$ |
| $S$ | $1.05 \times 10^{-2}$ | $1.76 \times 10^{-6}$ | $5.19 \times 10^{-8}$ | $1.57 \times 10^{-9}$ |
| $\Delta p_M$ | $3.59 \times 10^{-2}$ | $1.38 \times 10^{-7}$ | $4.29 \times 10^{-9}$ | $1.35 \times 10^{-10}$ |
| $E_{F,R}$ | $1.65 \times 10^{-3}$ | $2.31 \times 10^{-9}$ | $6.68 \times 10^{-11}$ | $1.99 \times 10^{-12}$ |
| $KE_{F,R}$ | $8.64 \times 10^{-3}$ | $6.38 \times 10^{-9}$ | $1.99 \times 10^{-10}$ | $5.77 \times 10^{-12}$ |

Ex vivo dual perfusion of the human placenta lobules was carried out at the University of Manchester under physiological conditions, and then perfusion fixed following infusion with Yasuaki contrast reagent [47]. The tissue was wax embedded for processing for *µ*CT imaging at the Diamond Light Source (Harwell campus, Oxfordshire). Synchrotron tomographic data was collected on the I13-2 beamline at the Diamond Light Source equipped with the robot autosampler following scanning protocol [28]. After reconstruction the *µ*CT data was segmented using an iterative human-in-the-loop deep learning process utlising a U-Net encoder-decoder architecture in volume segmantics [48].

### Problem Formulation

We follow a standard two-scale homogenisation approach to formulate a local cell problem from the Stokes equations, see e.g. [49]. The purpose of this is to calculate values of the effective permeability tensor on various local geometries, thereby to inform the choice of coefficient of the Darcy drag term in the Navier-Stokes-Darcy model (2).

To this end, guided by (2), we take the same characteristic scales as **S1 Table 1** with the exception of the characteristic length. Instead, we take the characteristic microscale length (pore scale) as 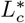 and the characteristic macroscale length (domain scale), 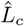, as a fictitious value such that 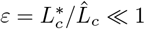. We emphasise that the intention here is not to obtain an upscaled macroscopic model, but rather to determine a suitable estimate for the effective permeability tensor for subsequent use in (2). In what follows, we hence focus on the local problem only.

We denote the local coordinate system by (*X*_1_, *X*_2_, *X*_3_) with unit basis vectors ***e***_*X*_ for *i* = 1, 2, 3. Differential operators taken with respect to this coordinate system. The unit cell is *Y* = [0, 1]^3^ with the flow domain *Y*_f_ ⊆ *Y* and solid domain *Y*_s_ ⊆ *Y* with *Y* = *Y* ∪ *Y* . Motivated by what follows, we also define *Ŷ* = [0, 1*/*2]^3^ with *Ŷ* and *Ŷ* the fluid and solid subdomains of *Ŷ*, respectively.

We have the periodic problems (forced periodic Stokes flow) on *Y*_*f*_

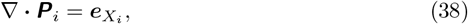

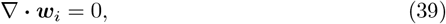

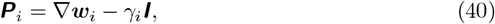

for *i* = 1, 2, 3, with ***w***_*i*_, *γ*_*i*_ periodic with period 1. The stress tensor here does not contain the (∇ ***w***_*i*_)^*T*^ for simplicity when presenting the derivation. However, the derivation remains the same and when solving the local system we will take the stress tensor in its symmetric form

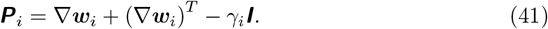

The domains for the unit cells are derived from *µ*CT scans, as described above. As the blocks are not periodic, they are not immediately suitable for the problem. Hence, to generate periodic blocks we instead generate a block from a *µ*CT scan which we take to represent the structure of *Ŷ* and reflect it 7 times on the *X*_1_, *X*_2_, *X*_3_ axes to form the periodic unit domain *Y* . The following analysis demonstrates that this step does not have to be done explicitly: instead a reduced problem can be solved on one eighth of the domain.

Due to the reflections, the periodic structure has reflectional symmetry. Via linearity of Stokes, we can show that this results in even-odd properties of the solution. To illustrate this, we consider *i* = 1 and write the equations out in component form

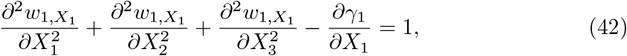

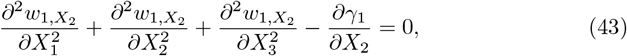

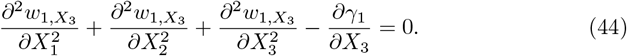

We see that by considering the substitutions 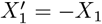 with the ansatz

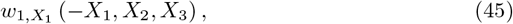

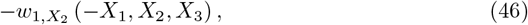

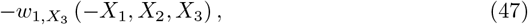

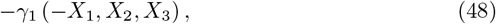

we get

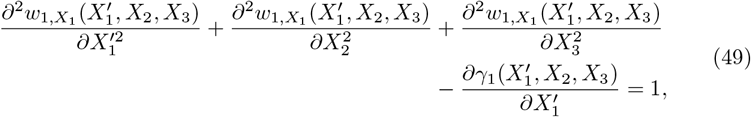

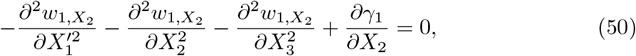

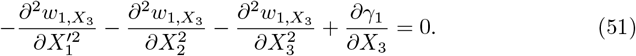

Hence, by uniqueness, the solution ***w***_1_, *γ*_1_ is such that 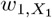 must be even in *X*_1_ and 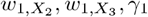 must be odd in *X*_1_. We can follow the same arguments for *X*_2_, *X*_3_ and *i* = 2, 3 to show for all local solutions

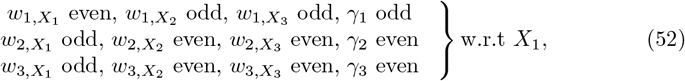

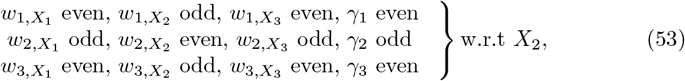

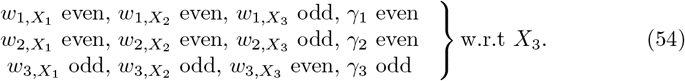

Using periodicity and (52)-(54), we have: for *i* = 1

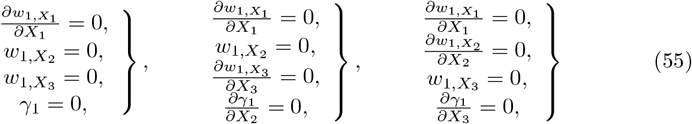

on *X*_1_ = 0, 1*/*2, *X*_2_ = 0, 1*/*2, *X*_3_ = 0, 1*/*2, respectively; for *i* = 2

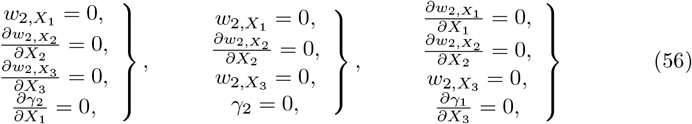

on *X*_1_ = 0, 1*/*2, *X*_2_ = 0, 1*/*2, *X*_3_ = 0, 1*/*2, respectively; for *i* = 3

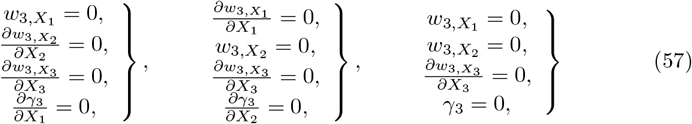

on *X*_1_ = 0, 1*/*2, *X*_2_ = 0, 1*/*2, *X*_3_ = 0, 1*/*2, respectively.

### Reduced Problem

From the above, we can define boundary conditions on n*X*_1_ = 0, 1*/*2, *X*_2_ = 0, 1*/*2, *X*_3_ = 0, 1*/*2 which allow us to reduce the problem onto *Ŷ*. For local problem *i*, the problem is

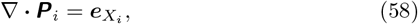

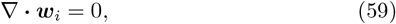

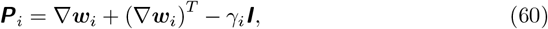

subject to

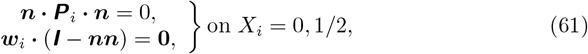

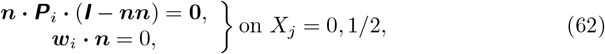

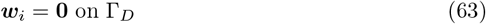

where *j* = 1, 2, 3, *j* ≠ *i*. Information on the numerical method used to solve this system can be found in **Numerical Method**.

### Permeability

The macroscopic permeability tensor, ***A***, is defined from the upscaled Darcy’s law

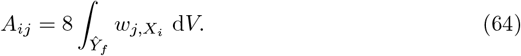

**S3 Fig 1.**
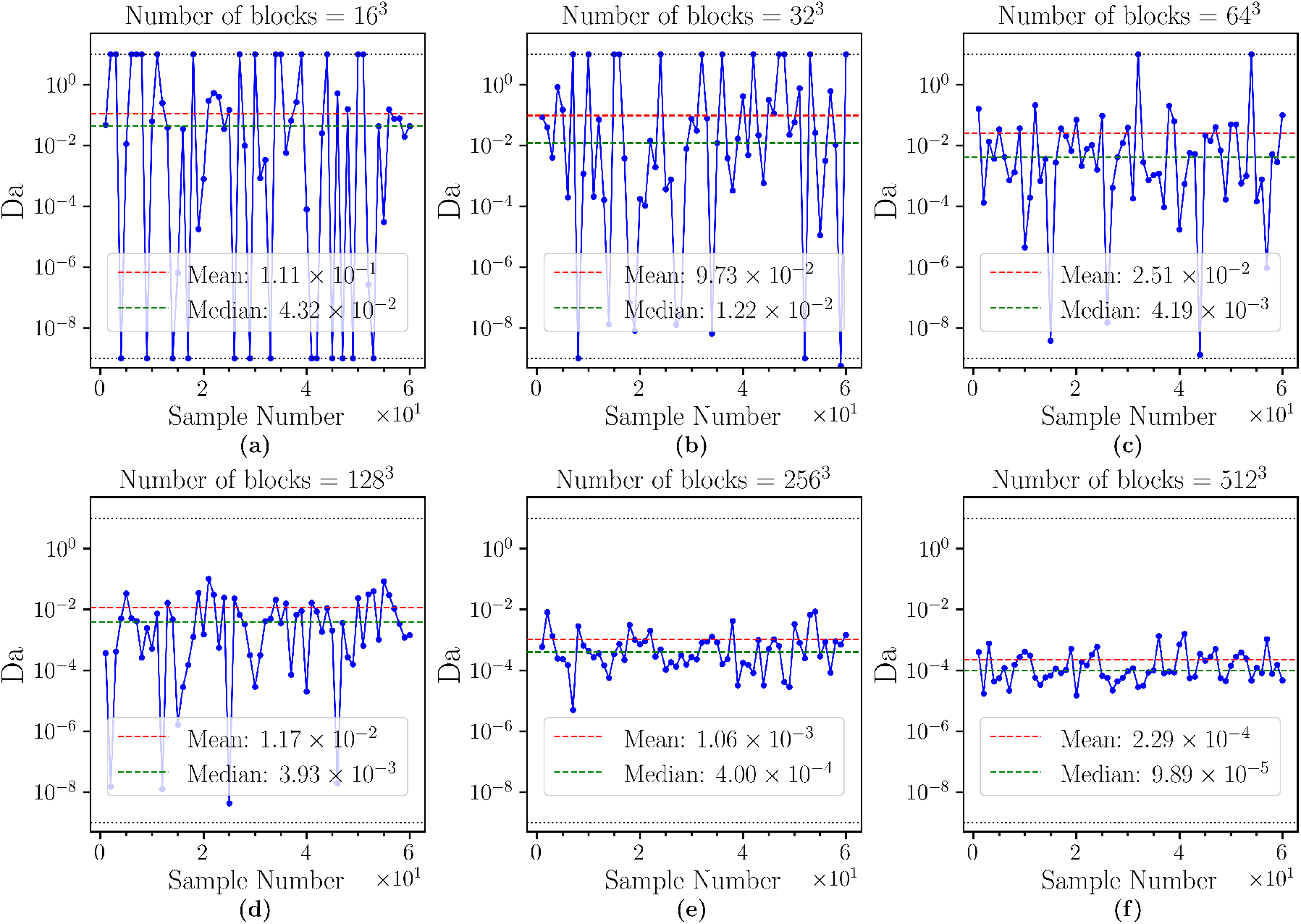
Permeability convergence with increasing data sample size. Plots of effective permeability for random fixed-size unit cube geometries.

We note that the factor of 8 is due to us solving the reduced problem on *Ŷ* rather than the unit cube *Y* .

To use this in the Navier-Stokes-Darcy model, we reduce this to an isotropic scalar by taking the Frobenius norm of *A*, i.e.,

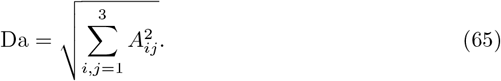

We remark that as the cell problem has been formulated from a non-dimensional problem, we maintain consistency with the organ-scale problem by denoting this effective permeability as Da. This is typically denoted *κ*^*^ when the upscaling is done in dimensional form.

In **S3 Fig 1**, we study the effect of sample size (i.e. number of blocks which comprises *Y*) on the calculated value of Da to assess whether convergence occurs under resolution refinement. Two situations can occur such that an ill-posed problem is formulated: if *Y* = *Y*_*f*_, there is no unique velocity solution; if *Y*_*f*_ = ∅, there is no flow domain. In these cases, we set the value of Da to be 10 and 10^−9^, respectively, for the purposes of visualisation. In the plots, when we calculate the mean and median of Da we ignore the values from the ill-posed cases.

**S4 Fig 1.**
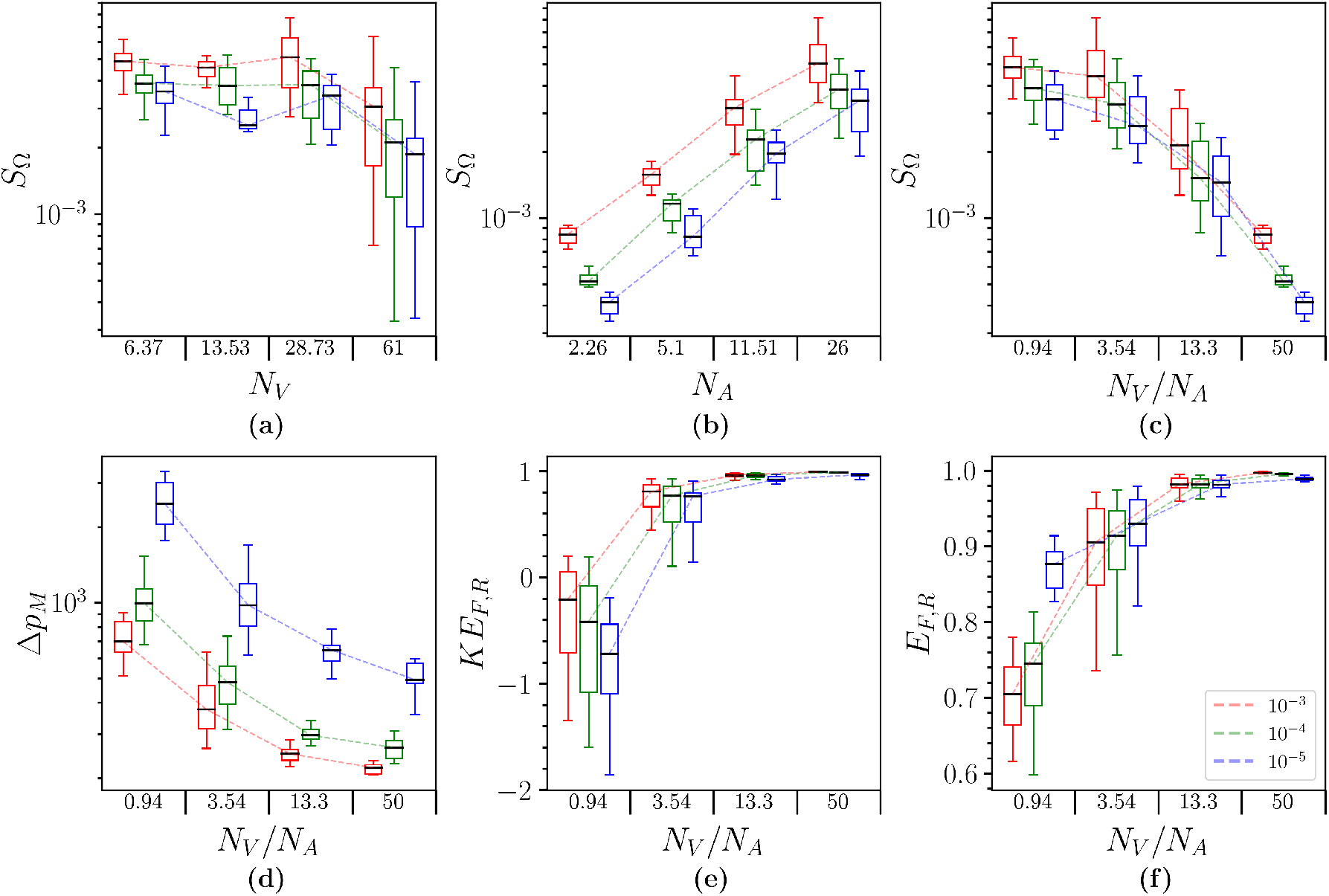
Placental haemodynamics box plots. Supporting box plot for Figure 7. The *x*-axis group number represents the upper bound for which the associated variable is part of that group. The legend in panel (c) indicates the Da value of each set of related box plots. For this data, 213 realisations were used with 72, 87, 54 realisations per Da in descending order that they appear in the legend. The non-dimensional number parameter ranges for this data are 4.54 *×* 10^2^ ≤ Re ≤ 1.36 *×* 10^3^.

## S4 Simulation Realisations and Supporting Plots

In the box plots presented here, **S4 Fig 1, S4 Fig 3** and **S4 Fig 4**, the *x* axis group value indicates the upper bound of the associated variable in that group, with the lower bound given by the left-adjacent group’s value (else 0 if it is the left-most group). For example, in **S4 Fig 4**(a) a data point of a particular *O*_Ω,up_ value belongs to the first group if the associated number of basal plate veins satisfies 0 ≤ *N*_*B*_ ≤ 2.56. Similarly, the second group is given by data points such that 2.56 *< N*_*B*_ ≤ 6.56.

In Figure 13(a,b), the corresponding fitted values are *a* = 1.01, *b* = 6.79 *×* 10^1^, *c* = 7.3 *×* 10^−1^ and *a* = 1.03, *b* = 5.34 *×* 10^1^, *c* = 6.17 *×* 10^−1^, respectively.

## S5 Kinetic Energy Flux Loss Ratio Approximate Law

In Figure 7(e), we plot *KE*_*F,R*_ against *N*_*V*_ */N*_*A*_. The dependence exhibits a characteristic pattern which is supported by analysing the consequences of mass conservation on *KE*_*F,R*_. First we note that in our flow simulations, arterial inflow is assumed to have a Poiseuille profile with the same (specified) maximum speed through flat, circular inlets. For the purposes of this analysis we make a further assumption that venous outflow also has a Poiseuille profile through the predefined flat, circular outlets, but with unknown maximum speed which is the same for each vein. Hence,

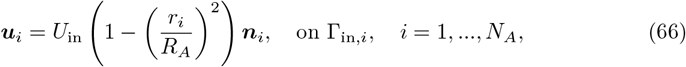

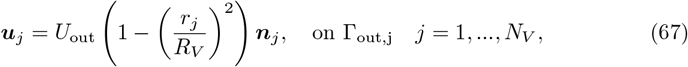

in which each *r*_*_ is a local radial coordinate in the plane perpendicular to the inward-pointing unit normal ***n***_*_ and centred on the origin of the circle defining the inlet or outlet face. *R*_*A*_, the artery radius, is assumed to be the same for all arteries. *R*_*V*_, the vein radius, will also be assumed to be the same for all veins when the approximations to *KE*_*F,R*_ are derived below: either 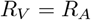 if all veins are assumed to be on either the basal plate or the septal walls or 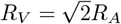 if all veins are assumed to be peripheral.

**S4 Fig 2.**
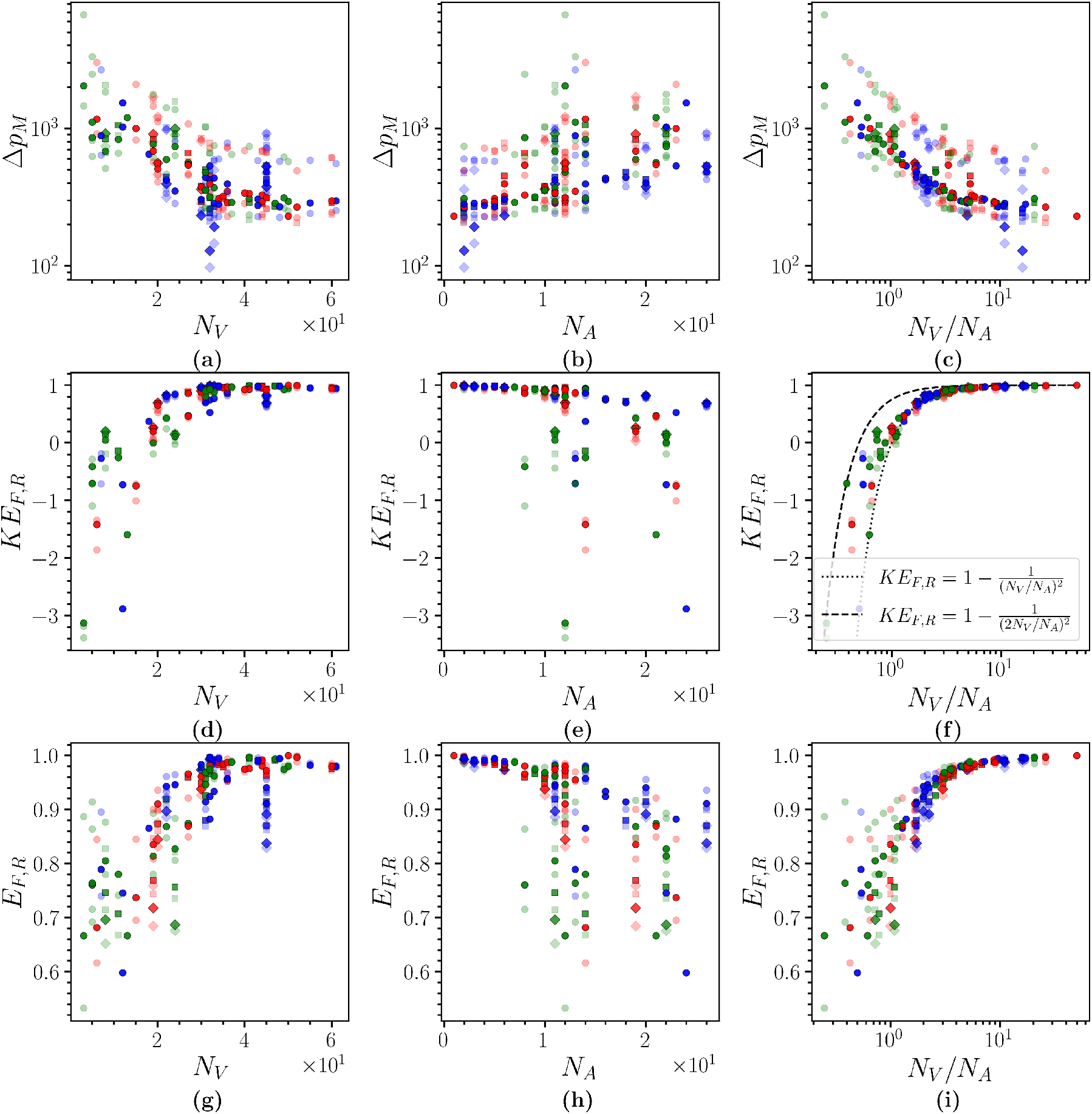
The variation of placental flow energy with vasculature. (a)–(c) Maximum mean arterial pressure drop, (d)–(f) kinetic energy flux loss ratio, (g)–(i) total energy flux loss ratio; against number of veins, arteries and vein-to-artery ratio, respectively. Each scatter point corresponds to a different random geometry and the colour indicates the cotyledon configuration it uses. The diamond, circle and square markers represent data with Da = 10^−3^, 10^−4^ (nominal), 10^−5^, respectively.

**S4 Fig 3.**
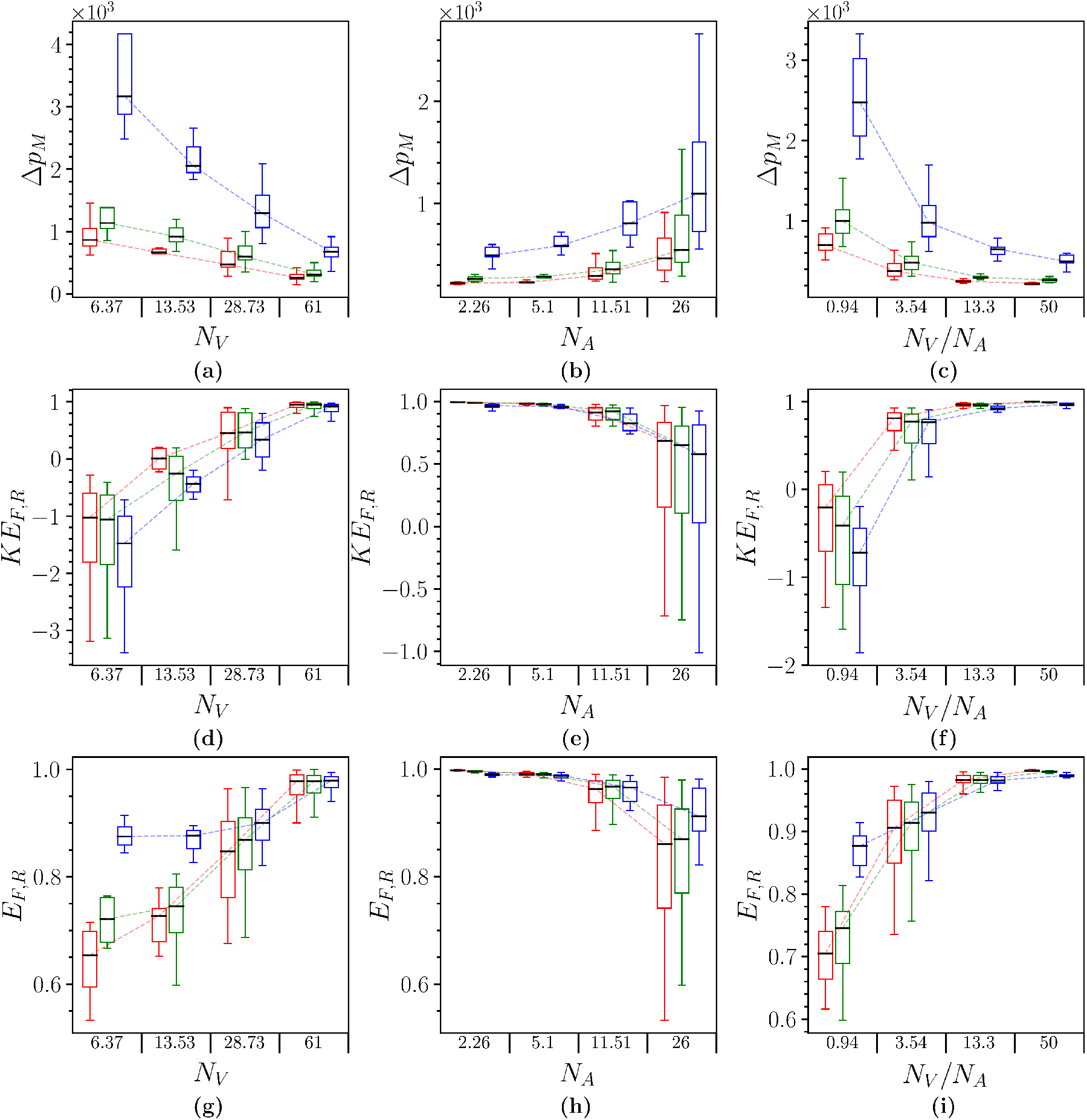
Placental flow energy box plot. Supporting box plot for **S4 Fig 2**. The *x*-axis group number represents the upper bound for which the associated variable is part of that group. The legend in panel (f) indicates the Da value of each set of related box plots. For this data, 213 realisations were used with 72, 87, 54 realisations per Da in descending order that they appear in the legend. The non-dimensional number parameter ranges for this data are 4.54 *×* 10^2^ ≤ Re ≤ 1.36 *×* 10^3^.

**S4 Fig 4.**
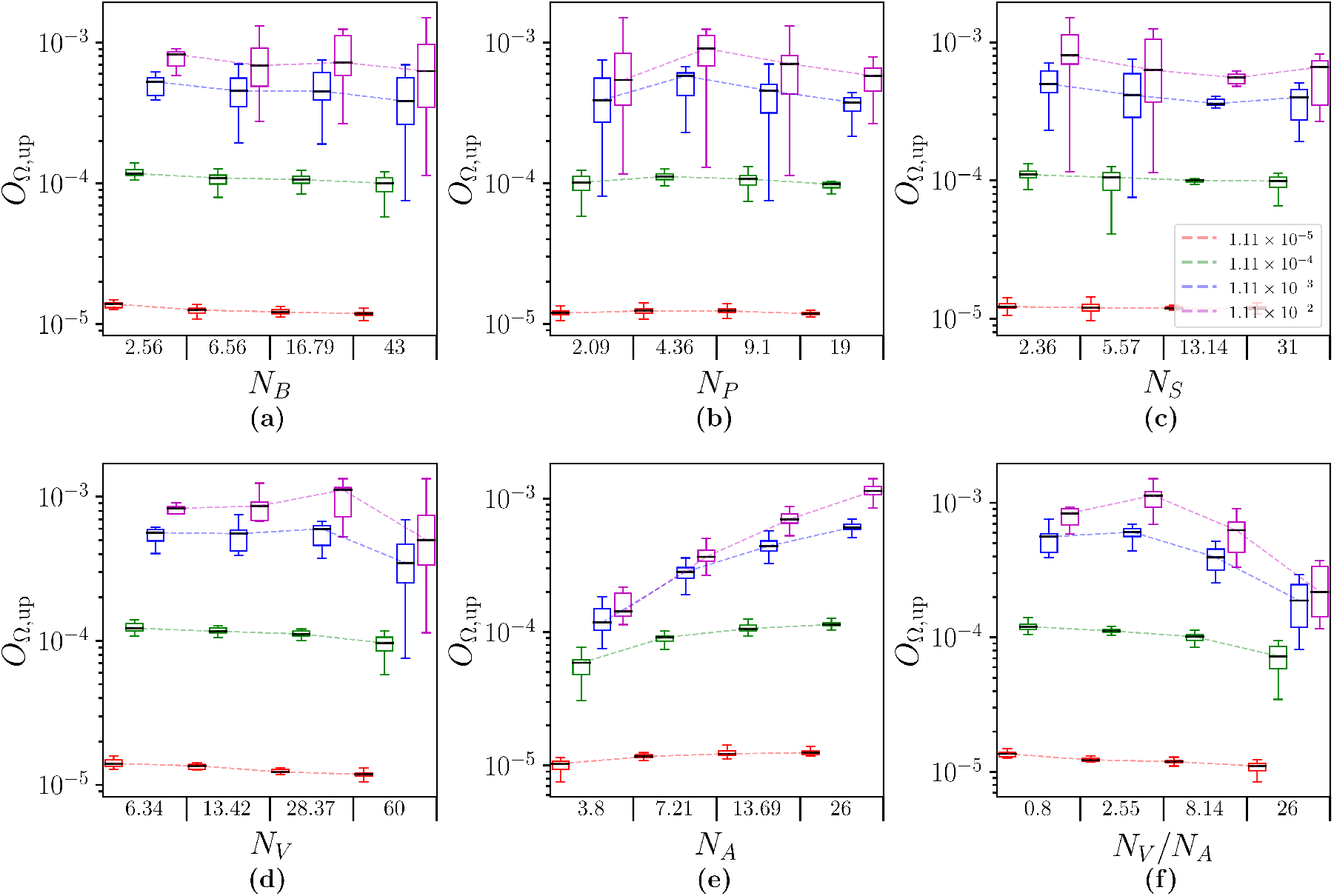
Oxygen uptake box plot. Supporting box plot for Figure 9. The *x*-axis group number represents the upper bound for which the associated variable is part of that group. The legend in panel (c) indicates the Dm value of each set of related box plots. For this data, 709 realisations were used with 175, 175, 184, 175 realisations per Dm in descending order that they appear in the legend. The non-dimensional number parameter ranges for this data are 9 *×* 10^5^ ≤ Pe ≤ 1.8 *×* 10^6^, 10^−5^ ≤ Da ≤ 10^−3^.

**S4 Fig 5.**
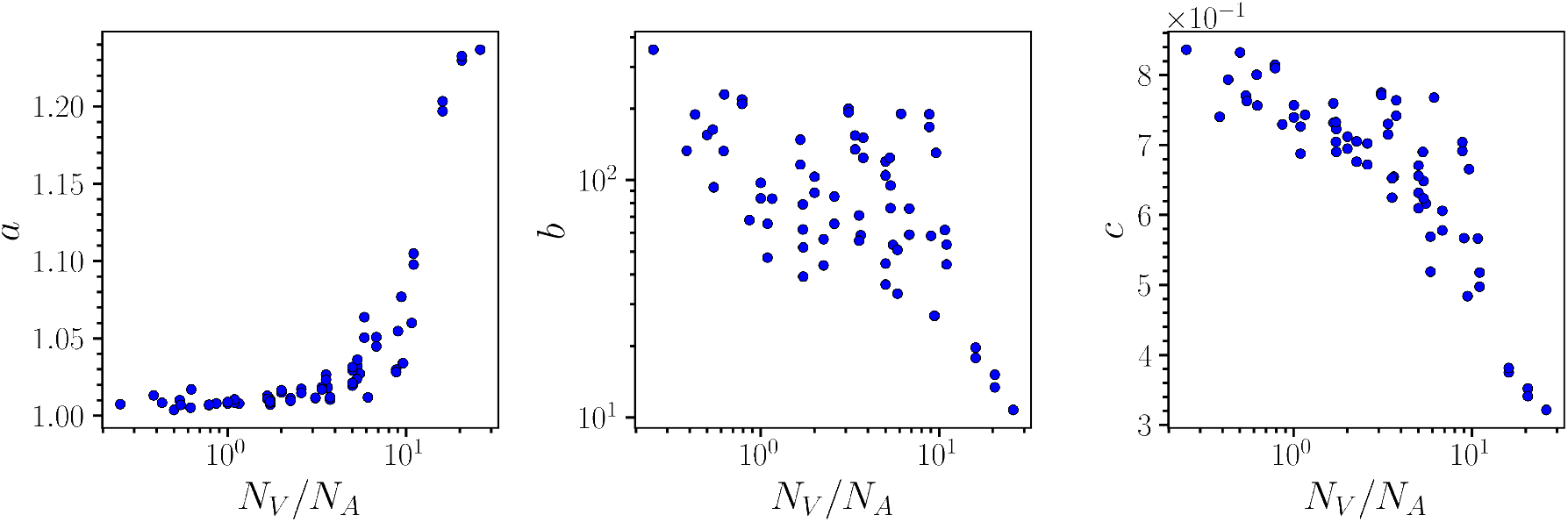
The variation of uptake function with vasculature. Fitting of the functional parameters for the uptake function ansatz specified in (36).

Now, by mass conservation,

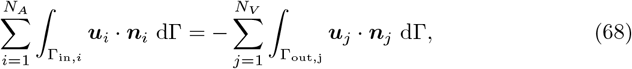

so substituting (66) and (67) gives, after integration,

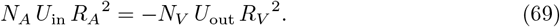

The inflow and outflow kinetic energy fluxes are given by

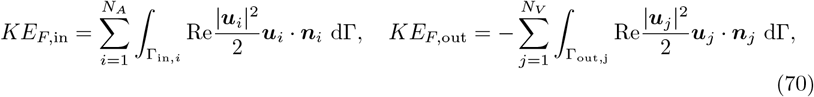

and substitution of (66) and (67), followed by integration gives

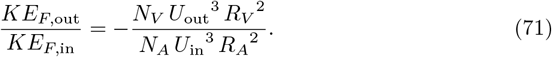

Combining this with (69) leads to

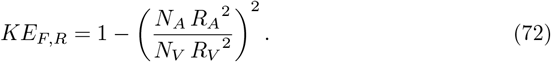

This analytical approximation is plotted in Figure 7 for *R*_*V*_ = *R*_*A*_ (dotted line – all veins are on the basal plate or septal walls) and 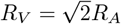 (dashed line – all veins are peripheral) and shows good agreement with the simulation data, especially for large *N*_*V*_ */N*_*A*_. At small *N*_*V*_ */N*_*A*_, the simulated results match more closely the dashed line due to the geometries having a higher proportion of the larger-radius peripheral veins.

## S6 Uptake in a half-space

To illustrate how solute exchange rate can be regulated by the source-sink separation distance, we consider the following idealised problem. Steady porous-medium flow occupies the half-space *y*^*^ *>* 0 and is driven by a point source with volume flux 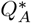 at the origin and two adjacent point sinks of strength 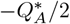 at 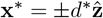. Elsewhere there is no flux through *y*^*^ = 0; the velocity field **u**^*^(**x**^*^) falls to zero in the far field. We assume axisymmetry with respect to the *z*-axis. The inlet concentration is 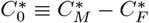 and we assume linear uptake kinetics of the solute distribution 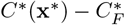 at rate 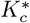. We scale lengths by *d*^*^, streamfunction by 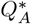, velocities by 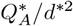, concentration by 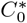, and time by 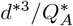. The solute exchange rate is 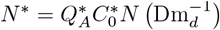 for some dimensionless function *N* (to be determined) of 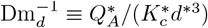. The dimensionless problem to be solved in the flow domain Ω is

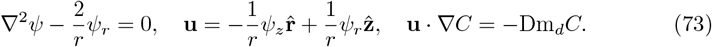

We seek the net solute exchange rate

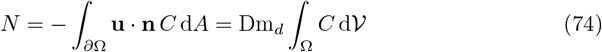

as a function of Dm_*d*_. Here **n** is the unit outward normal to *∂*Ω and fluxes across *∂*Ω are confined to the source and sinks. At the source, = 1 and ∫ **u n** · d = 1. At each sink, 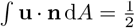 but the solute distribution must be calculated. This is done by tracking solute distribution along streamlines, which occupy stream-surfaces *ψ* = constant. Because of the symmetric arrangement of sinks, it is sufficient to consider *z >* 0.

From (73), following a particle in the flow, d*C/*d*t* = −Dm_*d*_*C* and d**x***/*d*t* = **u** on *ψ* = constant. Thus *C* = exp(−Dm_*d*_*t*) and so at the sink *C* = *C*_*s*_ = exp(−Dm_*d*_*t*_*s*_), where *t*_*s*_(*ψ*) is the transit time along the relevant streamsurface. *t*_*s*_ is computed by solving

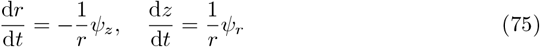

on *ψ* = constant, with *r* → 0, *z* → 0 as *t* → 0+, and *r* → 0, *z* → 1 as *t → t*_*s*_ − . Thus *C*_*s*_ = *C*_*s*_(*ψ*; Dm_*d*_).

The axisymmetric flow satisfying (73a) has streamfunction [24]

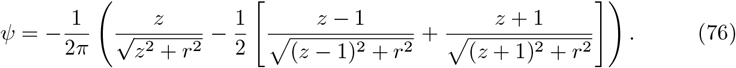

In the far field, with *r* ∼ *z* ≫ 1, (76) simplifies to

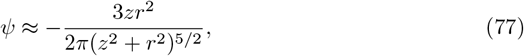

which is an axisymmetric dipole. To handle the singularities in (76), take hemispheres of radius *ρ* ≪ 1 around the source and the sink, incorporating these into *∂*Ω. Near the source, *ψ* is dominated by its first term and locally **u** ≈ − **n***/*(2*πρ*^2^). Near the sink at *z* = 1, *ψ* is dominated by its second term and locally **u** ≈ **n***/*(4*πρ*^2^). To evaluate the solute flux at the sink, we parametrize the surface of the hemisphere of radius *ρ* around it using *ψ*. The area element is d*A* = *πρ*^2^ sin *θ* d*θ* in spherical polar coordinates with *θ* = 0 pointing along 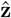, with 0 ≤ *θ* ≤ *π*. Locally *ψ* ≈ (−1 + cos *θ*)*/*(4*π*), so that d*A* = −*πρ*^2^(4*π* d*ψ*) for *ψ* = 0 to *ψ* = −1*/*(2*π*). Thus

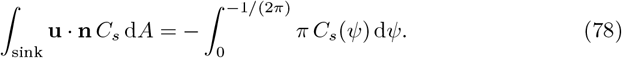

We now suppose Dm_*d*_ ≪ 1. Uptake is slow along streamlines, taking place predominantly in the far-field where the flow is described by (77). *C*_*s*_ ≈ 1 on the remaining streamlines that reach the sink without travelling far from the source; as these constitute the majority of streamlines, the integral in (78) approaches 1*/*2. However, we need the next correction in order to determine the rate of solute exchange. From (74),

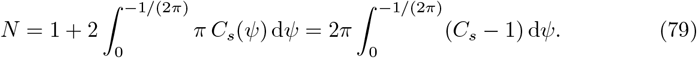

Treated as a function of position, *C* falls from unity near the source and sink, to zero in the very far field, over a region in which *ψ* is close to zero. (Streamlines with small *ψ* emerge from the source near the plane *z* = 0, and return to the sink along a thin region near the *z*-axis in *z >* 1. This slender thread penetrates the region in which *C* is otherwise close to unity.) In the far-field, *ψ* is approximated by (77), and the streamlines are self-similar. Thus, on a surface *ψ* = constant, we can write 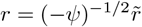 and 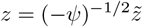 and 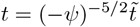. Then (75) and (77) become

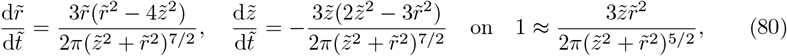

with 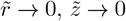 as 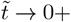 and 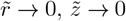 as 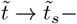. The integration of (80) provides the parametrization of the time of a path that originates in sector 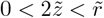 and returns to the origin in the sector 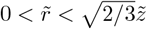. Thus the solute concentration at the sink, carried along streamlines that enter the far field, is

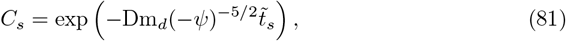

where 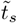 is a constant found to be approximately 0.0225 from numerical solution of (80). Now *C*_*s*_ ≈ 1 for Dm_*d*_(−*ψ*)^−5*/*2^ ≪ 1 (i.e.,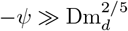). Thus (79) is determined by transport along the far-field streamlines, for which *ψ* is small. Setting 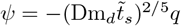, the exchange rate becomes

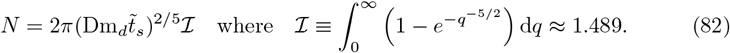

The dimensional solute exchange rate then satisfies

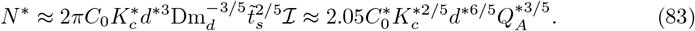

Equation (83) can be interpreted as a volume integral, with the concentration field being confined to a domain of dimensional radius 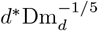 when the flow is strong. The approximation (83) is relevant provided 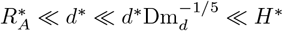, where 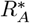 is the arterial radius at the source and *H*^*^ is the distance from the sources and sinks to the nearest boundary.

## Acknowledgments

This work was funded by the Wellcome Leap In Utero Programme (UoN authors: ‘Stillbirth: When Is Risk Low? (SWIRL)’; UoM authors: ‘Multi-modal studies to understand pregnancy and prevent stillbirth’). This work was carried out with the support of Diamond Light Source, instrument I13-2 (proposal MG32880). The authors thank Dr Michele Darrow, Dr Avery Pennington (Rosalind Franklin Institute), Dr Gowsihan Poologasundarampillai (University of Birmingham), Dr Paul Brownbill and Prof. Alexander Heazell (University of Manchester) for sharing the *µ*CT data.

## Data availability

The solver source code is available from Zenodo at DOI: 10.5281/zenodo.17580007. The simulation data comprising the figures in this manuscript, as well as the geometries and meshes used to specify the 3D sensitivity flow problems, are available from Zenodo at DOI: 10.5281/zenodo.18430960. Imaging data and associated segmentations are available upon request, please contact Michele Darrow at.

## Footnotes

1 This is done for performance markers associated with the 3D Navier-Stokes-Darcy equations.

## References

1. McLaughlin K, Scholten RR, Kingdom JC, Floras JS, Parker JD. Should Maternal Hemodynamics Guide Antihypertensive Therapy in Preeclampsia? Hypertension. 2018;71(4):550–556.

2. Valensise H, Farsetti D, Pometti F, Vasapollo B, Novelli GP, Lees C. The cardiac-fetal-placental unit: fetal umbilical vein flow rate is linked to the maternal cardiac profile in fetal growth restriction. American Journal of Obstetrics and Gynecology. 2023;228(2):222.e1–222.e12.

3. Lamont K, Scott NW, Jones GT, Bhattacharya S. Risk of recurrent stillbirth: systematic review and meta-analysis. BMJ. 2015;350:h3080.

4. Kolte AM, Westergaard D, Lidegaard Ø, Brunak S, Nielsen HS. Chance of live birth: a nationwide, registry-based cohort study. Human Reproduction. 2021;36(4):1065–1073.

5. Dellschaft NS, Hutchinson G, Shah S, Jones NW, Bradley C, Leach L, et al. The haemodynamics of the human placenta in utero. PLoS Biol. 2020;18(5):e3000676.

6. Knöfler M, Haider S, Saleh L, Pollheimer J, Gamage TKJB, James J. Human placenta and trophoblast development: key molecular mechanisms and model systems. Cell Mol Life Sci. 2019;76(18):3479–3496.

7. Campbell KA, Colacino JA, Puttabyatappa M, Dou JF, Elkin ER, Hammoud SS, et al. Placental cell type deconvolution reveals that cell proportions drive preeclampsia gene expression differences. Commun Biol. 2023;6(1):264.

8. Vermeulen RCW, Lambalk NB, Exalto N, Arts NFT. An anatomic basis for ultrasound images of the human placenta. American Journal of Obstetrics and Gynecology. 1985;153(7):806–810.

9. Wang Y. Vascular Biology of the Placenta, Second Edition. Colloquium Series on Integrated Systems Physiology: From Molecule to Function. 2017;9(3).

10. Kaufmann P, Miller RK, editors. Placental Vascularization and Blood Flow. Boston, MA: Springer US; 1988.

11. Schuhmann R, Stoz F, Maier M. In: Kaufmann P, Miller RK, editors. Histometric Investigations in Placentones (Materno-Fetal Circulation Units) of Human Placentae. Boston, MA: Springer US; 1988. p. 3–16.

12. Burton GJ, Woods AW, Jauniaux E, Kingdom JCP. Rheological and Physiological Consequences of Conversion of the Maternal Spiral Arteries for Uteroplacental Blood Flow during Human Pregnancy. Placenta. 2009;30(6):473–482.

13. Schiffer V, Evers L, De Haas S, Ghossein-Doha C, Al-Nasiry S, Spaanderman M. Spiral artery blood flow during pregnancy: a systematic review and meta-analysis. BMC Pregnancy Childbirth. 2020;20(1):680.

14. Brunelli R, Masselli G, Parasassi T, De Spirito M, Papi M, Perrone G, et al. Intervillous circulation in intra-uterine growth restriction. Correlation to fetal well being. Placenta. 2010;31(12):1051–1056.

15. James JL, Whitley GS, Cartwright JE. Shear stress and spiral artery remodelling: the effects of low shear stress on trophoblast-induced endothelial cell apoptosis. Cardiovascular Research. 2011;90(1):130–139.

16. Calis P, Vojtech L, Hladik F, Gravett MG. A review of ex vivo placental perfusion models: an underutilized but promising method to study maternal-fetal interactions. The Journal of Maternal-Fetal & Neonatal Medicine. 2022;35(25):8823–8835.

17. Capuani S, Guerreri M, Antonelli A, Bernardo S, Porpora MG, Giancotti A, et al. Diffusion and perfusion quantified by magnetic resonance imaging are markers of human placenta development in normal pregnancy. Placenta. 2017;58:33–39.

18. Moore R, Strachan B, Tyler D, Duncan K, Baker P, Worthington B, et al. In utero perfusing fraction maps in normal and growth restricted pregnancy measured using IVIM echo-planar MRI. Placenta. 2000;21(7):726–732.

19. Slator PJ, Hutter J, McCabe L, Gomes ADS, Price AN, Panagiotaki E, et al. Placenta microstructure and microcirculation imaging with diffusion MRI. Magnetic Resonance in Med. 2018;80(2):756–766.

20. Hutter J, Harteveld AA, Jackson LH, Franklin S, Bos C, Van Osch MJP, et al. Perfusion and apparent oxygenation in the human placenta (PERFOX). Magnetic Resonance in Med. 2020;83(2):549–560.

21. Hutchinson GJ, Blakey A, Jones N, Leach L, Dellschaft N, Houston P, et al. The effects of maternal flow on placental diffusion-weighted MRI and intravoxel incoherent motion parameters. Magnetic Resonance in Med. 2025;93(4):1629–1641.

22. Jensen OE, Chernyavsky IL. Blood Flow and Transport in the Human Placenta. Annu Rev Fluid Mech. 2019;51(1):25–47.

23. Erian FF, Corrsin S, Davis SH. Maternal, placental blood flow: A model with velocity-dependent permeability. Journal of Biomechanics. 1977;10(11):807–814.

24. Chernyavsky IL, Jensen OE, Leach L. A Mathematical Model of Intervillous Blood Flow in the Human Placentone. Placenta. 2010;31(1):44–52.

25. Lecarpentier E, Bhatt M, Bertin GI, Deloison B, Salomon LJ, Deloron P, et al. Computational Fluid Dynamic Simulations of Maternal Circulation: Wall Shear Stress in the Human Placenta and Its Biological Implications. PLoS ONE. 2016;11(1):e0147262.

26. Saghian R, James JL, Tawhai MH, Collins SL, Clark AR. Association of Placental Jets and Mega-Jets With Reduced Villous Density. Journal of Biomechanical Engineering. 2017;139(5):051001.

27. Baergen RN, Burton GJ, Kaplan CG, editors. Benirschke’s Pathology of the Human Placenta. Cham: Springer International Publishing; 2022. Available from: https://link.springer.com/10.1007/978-3-030-84725-8.

28. Tun WM, Poologasundarampillai G, Bischof H, Nye G, King ONF, Basham M, et al. A massively multi-scale approach to characterizing tissue architecture by synchrotron micro-CT applied to the human placenta. J R Soc Interface. 2021;18(179):20210140.

29. De Berg M, Cheong O, Van Kreveld M, Overmars M. Computational Geometry: Algorithms and Applications. Berlin, Heidelberg: Springer Berlin Heidelberg; 2008.

30. Blakey AM. Placental haemodynamics: a computational study of maternal blood flow and oxygen transport in the human placenta [PhD thesis]. University of Nottingham. Nottingham, Nottinghamshire; 2024.

31. Mekler T, Plitman Mayo R, Weissmann J, Marom G. Impact of tissue porosity and asymmetry on the oxygen uptake of the human placenta: A numerical study. Placenta. 2022;129:15–22.

32. Carson J, Warrander L, Johnstone E, Van Loon R. Personalising cardiovascular network models in pregnancy: A two-tiered parameter estimation approach. Numer Methods Biomed Eng. 2021;37(11):e3267.

33. Kurjak A, Dudenhausen JW, Hafner T, Kupesic S, Latin V, Kos M. Intervillous circulation in all three trimesters of normal pregnancy assessed by color Doppler. J Perinat Med. 1997;25:373–380.

34. Collins SL, Stevenson GN, Noble JA, Impey L. Developmental changes in spiral artery blood flow in the human placenta observed with colour Doppler ultrasonography. Placenta. 2012;33(10):782–787.

35. Allaire G. Homogenization and Two-Scale Convergence. SIAM J Math Anal. 1992;23(6):1482–1518.

36. Pavliotis GA, Stuart AM. Multiscale Methods. Texts Applied in Mathematics. New York, NY: Springer New York; 2008. Available from: http://link.springer.com/10.1007/978-0-387-73829-1.

37. Chernyavsky IL, Leach L, Dryden IL, Jensen OE. Transport in the placenta: homogenizing haemodynamics in a disordered medium. Phil Trans R Soc A. 2011;369(1954):4162–4182.

38. Erlich A, Pearce P, Mayo RP, Jensen OE, Chernyavsky IL. Physical and geometric determinants of transport in fetoplacental microvascular networks. Science Advances. 2019;5(4):eaav6326.

39. Cockburn B, Karniadakis GE, Shu CW. Discontinuous Galerkin Methods. Berlin, Heidelberg: Springer Berlin Heidelberg; 2000.

40. Shahbazi K, Fischer PF, Ethier CR. A high-order discontinuous Galerkin method for the unsteady incompressible Navier–Stokes equations. Journal of Computational Physics. 2007;222(1):391–407.

41. Jaffre J, Johnson C, Szepessy A. Convergence of the Discontinuous Galerkin Finite Element Method for Hyperbolic Conservation Laws. Math Models Methods Appl Sci. 1995;05(03):367–386.

42. Hartmann R, Houston P. Adaptive Discontinuous Galerkin Finite Element Methods for the Compressible Euler Equations. Journal of Computational Physics. 2002;183(2):508–532.

43. Forti D, Dedè L. Semi-implicit BDF time discretization of the Navier–Stokes equations with VMS-LES modeling in a High Performance Computing framework. Computers & Fluids. 2015;117:168–182.

44. Cliffe KA, Hall EJC, Houston P. Adaptive Discontinuous Galerkin Methods for Eigenvalue Problems Arising in Incompressible Fluid Flows. SIAM Journal on Scientific Computing. 2010;31(6):4607–4632.

45. Du Q, Faber V, Gunzburger M. Centroidal Voronoi Tessellations: Applications and Algorithms. SIAM Review. 1999;41(4):637–676.

46. MATLAB version 23.2.0.2515942 (R2023b) Update 7; 2023.

47. Tokudome Y, Poologasundarampillai G, Tachibana K, Murata H, Naylor AJ, Yoneyama A, et al. Curable Layered Double Hydroxide Nanoparticles-Based Perfusion Contrast Agents for X-Ray Computed Tomography Imaging of Vascular Structures. Advanced NanoBiomed Research. 2022;2(2):2100123.

48. Bellos D, Basham M, et al. Volume Segmantics: A Python Package for Semantic Segmentation of Volumetric Data Using Pre-trained PyTorch Deep Learning Models. Journal of Open Source Software. 2022;7(78):4691.

49. Sanchez-Palencia E. Non-Homogeneous Media and Vibration Theory. Berlin, Heidelberg: Springer Berlin Heidelberg; 1980.

